# Mechanochemical Energy Landscapes Under Force: Catch–Slip Bonds in T-Cell Activation

**DOI:** 10.1101/2025.11.15.688643

**Authors:** Yogesh Bali, Alan D. Rendall, Wolfgang Quapp

## Abstract

Traditional theories explain T-cell activation by antigen concentration, binding affinity, and dissociation rate, but they fail to account for its extraordinary sensitivity. Recent cryo-electron microscopy and molecular-dynamics studies show that mechanical force transmitted through the T-cell receptor, the peptide–major histocompatibility complex, and the CD3 signaling module can enhance this sensitivity and thereby promote T-cell activation. We present a minimal energy-landscape model showing how force transmission reshapes the activation barrier and bond lifetime. Mathematically, we prove a sharp force cutoff *F*_*c*_ = *D*_0_/(2*d*_0_) (maximum binding depth *D*_0_, optimal distance *d*_0_) above which no bound state exists. At or below *F*_*c*_, the system admits at most two stationary points: a bound minimum and a saddle that coalesce in a fold (saddle-node) at the bond-breaking point. Stable binding occurs only on the near-separation branch *d < d*_0_ ln 2 (smaller receptor-ligand separation). Numerically, we tracked these stationary points as the force varied, computed the force-dependent barrier Δ*V* _*f*_, and mapped bond lifetimes using Kramers/Langer theory. We found that agonists and weak agonists follow a catch–slip pattern; barriers and lifetimes peak at intermediate forces, whereas antagonists exhibit slip, with barriers and lifetimes decreasing as force increases. Our framework provides a testable mechanochemical link between force, barriers, and lifetimes in TCR signaling, as well as a path to refine predictions with targeted measurements for T-cell activation under force.

## 1 Background

The interaction of T cell receptors (TCRs) with peptides presented by major histocompatibility complex (MHC) proteins is central to cellular immunity. Classical views emphasized receptor–ligand affinity and dissociation kinetics as the primary drivers of T-cell activation. However, TCR-pMHC binding represents one of the weakest protein-protein interactions, with affinities ranging from 10^−4^ to 10^−6^ M-approximately 1000 times weaker than antigen-antibody interactions (10^−6^ to 10^−10^ M) [1, 2]. Despite this low affinity, TCRs exhibit extraordinary sensitivity and specificity, capable of recognizing a single foreign pMHC among numerous self-antigens in antigen-presenting cells (APCs) [3], and in some settings even responding to a single pMHC [4]. The mechanism behind this high sensitivity, despite weak binding, has been a major focus of research [5].

It was hypothesized that mechanical forces modulate TCR-pMHC sensitivity and specificity by inducing conformational changes in the receptor-ligand complex, thereby improving the TCR’s ability to distinguish between agonist and non-agonist peptides [6]. Subsequent work indicated that the *αβ* TCR operates as a mechanosensor, detecting mechanical forces to regulate activation [7].

Kim et al. [8] provided some of the earliest compelling evidence supporting the role of TCRs as mechanosensors. Using optical tweezers (OT), biomembrane force probes (BFP) and nuclear magnetic resonance (NMR), they reported orientation-dependent engagement and anisotropic force sensitivity of the TCR–CD3 complex, and showed that applying tangential forces in the tens of piconewtons (∼10–50 pN) to non-agonist antigens enabled T cell activation, highlighting the critical role of mechanical forces in TCR activation.

Similarly, Lee et al. [9] demonstrated the role of mechanical forces for TCR activation by using artificial APCs expressing elongated anti-CD3 antigens. T cells bound to these artificial APCs failed to activate in the absence of external mechanical forces. However, when such forces were applied, effective T cell activation occurred, further reinforcing the notion that mechanical transmission through the TCR complex is essential for initiating signaling.

Further studies showed that applying a force of approximately 10 pN to the bond between a TCR and a specific pMHC can prolong bond lifetime. Activating peptides were observed to form longer-lived bonds with TCRs under mechanical force, while antagonist peptides failed to maintain the bond under similar conditions [10, 11]. This finding underscores the role of mechanical forces in distinguishing between antigens. Furthermore, fluorescence-enabled biomembrane force probe (fBFP) measurements have shown that agonist TCR-pMHC interactions exhibit catch-bond behavior; bond lifetimes increase under applied force thereby allowing sufficient time for productive T cell activation; in contrast, antagonist ligands form slip bonds where bond lifetimes monotonically decrease with force, limiting signaling [12]. This phenomenon, where the lifetimes of the bonds initially increase under applied force, reach a peak at an optimal force level, and then decline as the force continues to increase, is referred to as a catch bond [10, 13].

In T cells, catch bonds typically peak around 10–15 pN [14]; by contrast, slip bonds weaken as force rises [15, 16]. Since catch bonds are strengthened by moderate load, they extend the contact time between the TCR and the ligand, providing additional time for signaling to occur [17, 18]. This prolonged interaction facilitates time-dependent processes, such as protein segregation and structural changes in the TCR driven by allostery, where binding at one site influences other sites on the protein [19–22]. In fact, this mechanical force is thus pivotal not only for TCR triggering but also for T cell development in the thymus, including positive selection [23–25].

Since multiple studies indicate that applying force to TCR-pMHC prolongs bond lifetimes and helps in T-cell activation, several mechanisms have been proposed to explain how load is converted into productive structural changes at the receptor. In T cell signaling, a key early event is phosphorylation of the tyrosines within immunoreceptor tyrosine-based activation motifs (ITAMs) [26, 27]. These ITAMs are presented on CD3 *ε*; in resting T cells, the cytoplasmic tails of the TCR complex—particularly CD3 *ε*—are associated with acidic lipids and can be partially embedded in the inner leaflet of the plasma membrane [28]. It was assumed that when TCRs engage antigens, local membrane bending generates steric and lateral pressure, driving a conformational change, potentially pushing the cytoplasmic tails out of the membrane. This exposes the ITAMs, allowing lymphocyte-specific protein tyrosine kinase (Lck) to phosphorylate them and trigger T-cell activation [29]. Changes in membrane curvature also affect how lipids are packed, which can further influence TCR signaling dynamics [30, 31]. Furthermore, the extent of these conformational changes is believed to correlate with the potency of antigens, with stronger antigens inducing the necessary conformational changes more effectively [32].

### TCR structure and how conformational change occurs

The TCR is a heterodimeric glycoprotein composed of *α* and *β* polypeptide chains (TCR-*αβ*). Each chain has a variable (V) region responsible for antigen recognition and a constant (C) region for structural support. The V region comprises three complementarity determining regions (CDRs), with CDR3 being central to peptide antigen binding. Because the TCR lacks intrinsic signaling capacity, it depends on the CD3 complex for signal transduction [33, 34]. The CD3 complex is composed of six non-polymorphic chains, organized into CD3*γ*-CD3*ε* and CD3*δ*-CD3*ε* heterodimers, along with a CD3*ζ ζ* homodimer, all arranged in specific orientations relative to the TCR [35, 36]. A labeled view of the complete TCR–pMHC–CD3 assembly is shown in Fig. 1.

**Fig. 1.**
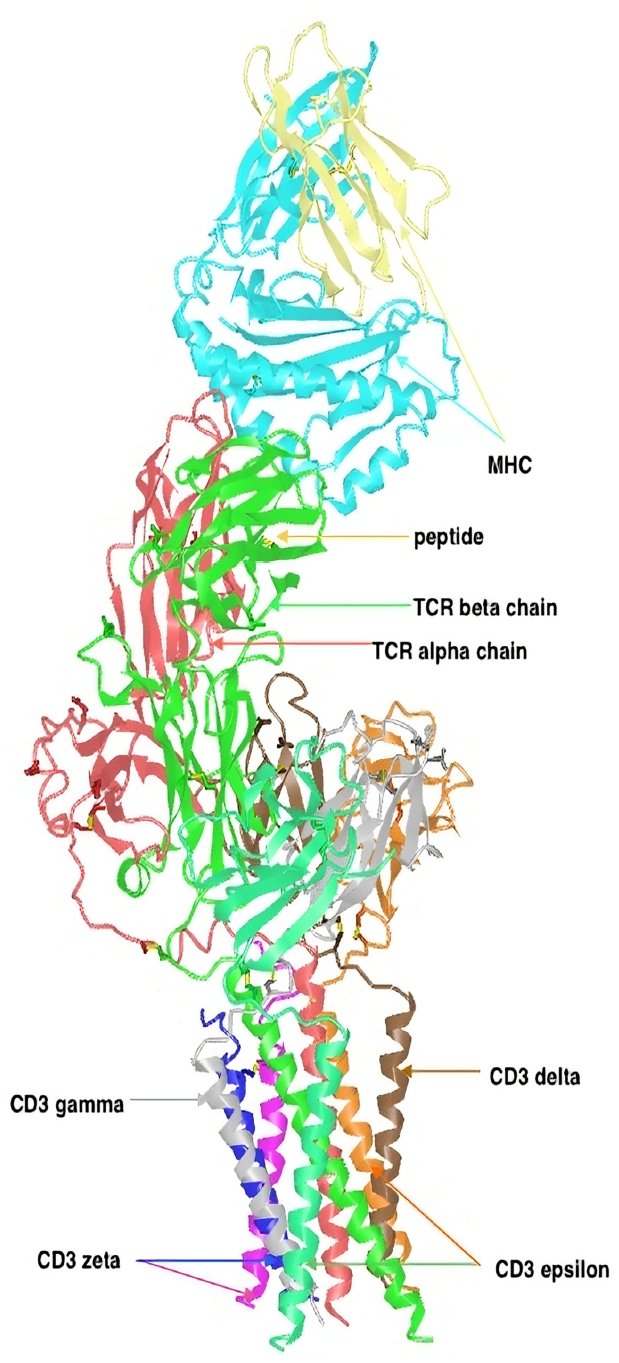
Labeled view of the TCR–pMHC–CD3 assembly. The peptide (cyan) lies in the MHC groove (yellow) and is recognized by TCR *α* (green) and TCR *β* (red). CD3 subunits are colored as: *ε* (orange), *δ* (brown), *γ* (gray), and *ζ* (purple/blue). Rendered in iCn3D from cryo-EM coordinates (e.g., PDB 8ES8/8ES9); see Saotome *et al*. [45] for experimental details.

Recent advancements in cryo-electron microscopy (cryo-EM) and molecular dynamics (MD) simulations have provided critical insights into the structural organization and membrane alignment of the TCR–CD3 complex [37–40]. These studies focus on delineating interactions between TCR *αβ* and CD3 subunits and on characterizing potential conformational changes in the extracellular (EC) domain of TCR *αβ* relative to the membrane. Upon pMHC engagement, these rearrangements couple to the phosphorylation-driven sequence—membrane disengagement of CD3 tails, ITAM exposure, and Lck access—outlined in 1, thereby linking geometry to early biochemical signaling.

In 2019, single-particle cryo-EM resolved the full-length TCR–CD3 complex and mapped the chain–chain interfaces. The TCR*α* chain interacts with CD3*δ* at the DE loop (the connector between *β*-strands D and E in the Ig-like V*α* domain) and connecting-peptide (CP) region, with additional stabilization from CD3*ζ*, while the TCR*β* chain interfaces with CD3*γ* and CD3*ε* mainly in their extracellular and membrane-proximal segments [40]. Subsequent cryo-EM and NMR studies refined this picture: the TCR*α* DE loop engages the CD3*εδ* module, whereas the TCR*β* CC^′^ loop (between *β*-strands C and C^′^ in V*β*) contacts both CD3*εδ* and CD3*εγ*; by contrast, the TCR C*β* FG loop (between *β*-strands F and G in the constant *β* domain) shows only limited interaction with CD3*εγ*, consistent with cooperative rather than uniformly strong binding across the assembly [41, 42].

MD simulations extended these structural findings, revealing broader and more persistent contacts involving the TCR C*β* FG loop (the connector between *β*-strands F and G in the constant *β* domain) and the TCR*α* variable domain (V*α*) than initially appreciated [39]. Notably, comparisons of TCR–CD3 alone with the TCR–CD3–pMHC complex show a marked reduction in TCR–CD3 contacts upon pMHC engagement. This rearrangement is proposed to constrain the spatial freedom of CD3 subunits and to expose or activate signaling motifs within CD3 cytoplasmic tails, thereby initiating downstream signaling and sharpening specificity [20]. Further, the TCR extracellular domain adopts a bent conformation (≈ 104^°^ tilt) in the resting state and transitions to a more extended orientation (≈ 150^°^) upon pMHC engagement (see Fig. 2). This reorganization reorients the TCR relative to CD3, facilitating signal transmission [38, 43]. In addition, the CD3 *ε* chains exhibit increased flexibility and motion in the presence of pMHC, exploring regions inaccessible in the unbound state [39]. This conformational flexibility supports catch bond behavior, akin to selectin catch–slip dynamics, may enhance the TCR’s ability to engage diverse pMHC molecules and transmit activation signals [7, 44].

**Fig. 2.**
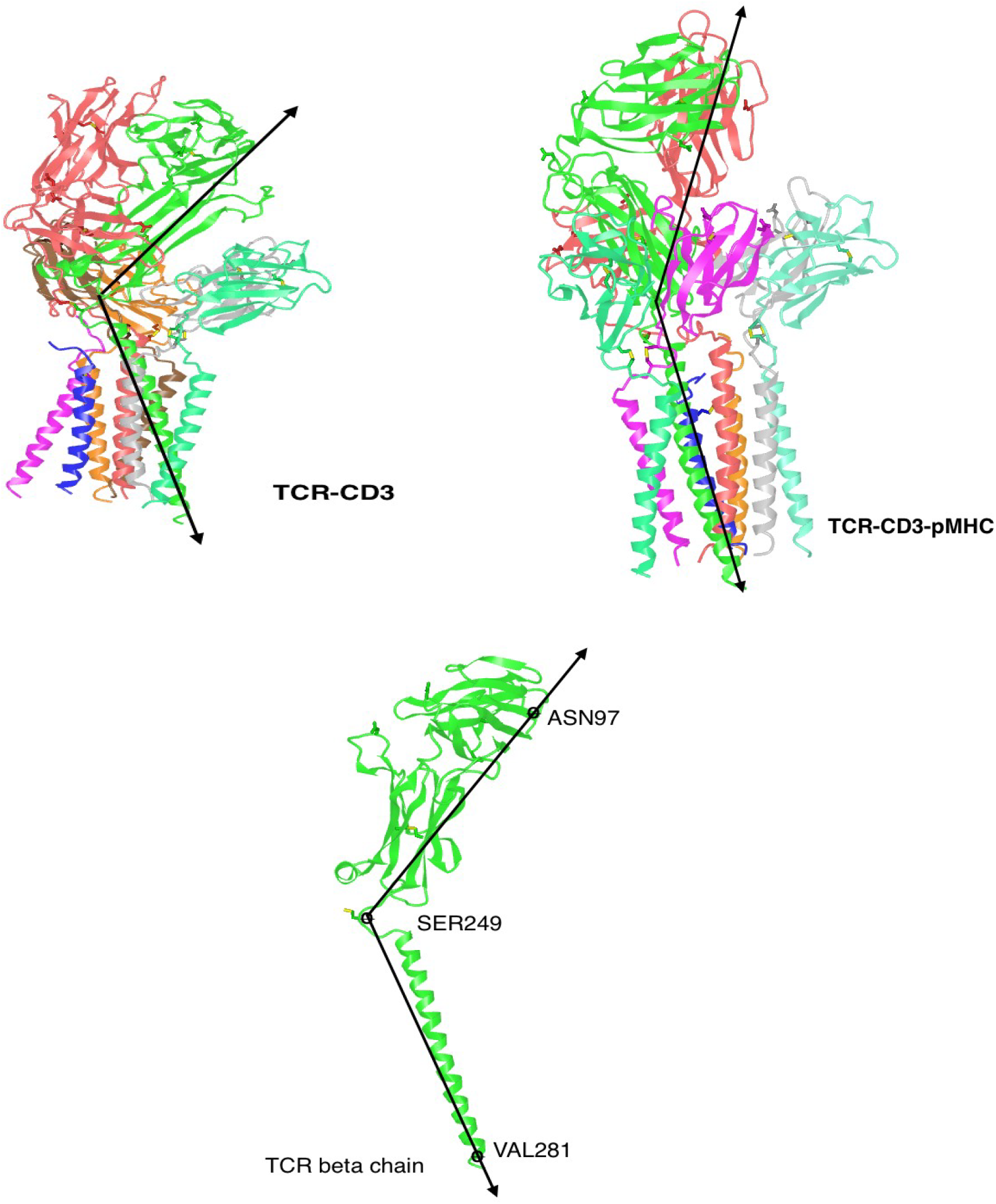
Tilt-angle comparison and definition across the TCR–CD3 assembly. Left: TCR–CD3 baseline (no pMHC). Right: pMHC-engaged state *(pMHC not shown)* with increased TCR*β* tilt; same viewing orientation as the left panel for direct comparison. Bottom: residue-based definition of the TCR*β* tilt angle *θ* using ASN_97_, SER_249_, and VAL_281_. Other chains are colored distinctly for clarity (e.g., TCR*α* in red/magenta; CD3 subunits in varied colors).

## 2 Model Development

Mechanochemistry is an emerging field focused on studying how mechanical forces initiate or modulate chemical reactions [46, 47]. Theoretically, applying external forces perturbs the potential energy surface (PES) of a molecular system, leading to shifts in the locations of minima and saddle points (SPs) [48]. These changes influence the energy barriers separating minima and SPs, thereby altering the system’s reaction pathways. The extent and nature of these modifications depend on both the strength and direction of the applied force [48], resulting in either an increase or decrease in chemical reactivity.

The catch-slip bond phenomenon is a prominent example of how mechanical forces can modulate chemical reactivity. In our model, this behavior can be understood as arising from force-induced alterations of the potential energy surface (PES), leading to two effective regimes [49]. At low to intermediate forces, load can stabilize the bound configuration relative to the transition state, consistent with a catch-like increase in bond lifetime; at higher forces, the same load promotes barrier lowering and bond rupture [50, 51], in line with slip-like behavior reported for TCR catch bonds.

To capture the dynamics of catch-slip bonds, simplified, coarse-grained models of the PES have been developed. These models provide a clear representation of how mechanical forces affect PES and, in turn, influence bond stability and reaction mechanisms. By incorporating these models, researchers can better understand the interplay among force, energy landscapes, and chemical reactivity, providing mechanistic insight into mechanochemistry [52].

As indicated by SMD simulations, as well as studies employing OT [53], BFP, magnetic tweezers [54], and cryo-EM techniques [39], the TCR-pMHC interactions under force are associated with the formation of catch-slip bonds. These studies revealed that the TCR-pMHC complex undergoes elastic expansion as it transitions from the bound state to the transition state. This process is accompanied by structural changes, including hinge rotations and the unfolding of interdomain connections [54]. Specifically, the angle between the TCR *β* trans membrane domain (TM) and the extracellular domain (EC) increases upon binding with pMHC [38, 55], providing the mechanical coupling that links force application to biochemical signaling.

Newton trajectory theory serves as a foundational framework for understanding the biochemical mechanisms behind slip and catch bond behavior [49, 56]. A key concept in analyzing molecular systems and their stable or unstable states is PES, which maps a molecule’s potential energy as a function of its atomic positions. This creates a multidimensional landscape where each point represents a unique molecular configuration. Within this landscape, local minima correspond to stable states where the system is in equilibrium, while saddle points represent transition states that molecules must cross to move between stable configurations [57].

The application of an external force to a bond, particularly in systems exhibiting catch and slip bonds, alters both the stable bond configuration (local energy minimum) and the transition state (saddle point). This force-induced modification shifts the positions of these critical points within the configuration space, thereby influencing the bond’s stability and dynamic behavior [58, 59]. To effectively analyze these changes, a multidimensional understanding of the PES is required. This complexity is often simplified using coarse-grained models, which focus on large-scale molecular motions to capture the system’s essential dynamics. Such models provide critical insights into how force-driven alterations in the PES dictate the formation and stability of catch and slip bonds.

The potential energy landscape of a molecular system under external force, denoted *V* _*f*_ (**x**), is commonly modeled as [49, 56]:

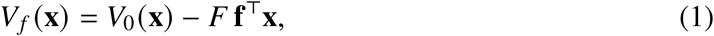

where:

- *V*_0_(**x**) is the intrinsic (unloaded) PES of the system.
- The term *F* **f**^⊤^**x** represents the effect of a constant force *F* applied along the unit direction **f** (superscript ^⊤^ denotes transpose) [60].

This framework captures how mechanical loading reshapes the energy landscape and moves stationary points (minima and saddle points) in configuration space [57, 61]. The barrier between a saddle **x**_sp_ and a local minimum **x**_min_ on the loaded surface is

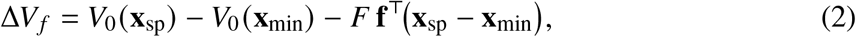

i.e., an intrinsic term minus a force-dependent energy shift.

At stationary points **x**_*c*_, the gradient of the effective PES vanishes, which leads to

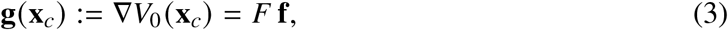

where **g**(**x**_*c*_) is the gradient of the intrinsic PES [48]. Curves satisfying this relation for varying *F* are Newton trajectories (NTs). For any **x**_*c*_, the corresponding force magnitude is

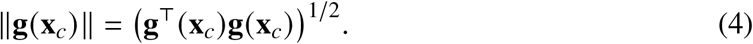

These trajectories describe how molecular configurations evolve with applied force. To describe the motion of any critical point **x**_*c*_, we formulate a corresponding differential equation.:

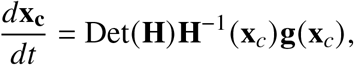

Here, **H**^−1^ denotes the inverse of the Hessian matrix associated with the intrinsic (PES), and Det(**H**) represents the determinant of this matrix. The variable *t* is a parameter along the Newton trajectory [62, 63]. To suppress Hessian singularities, a regularized matrix **A** is defined as:

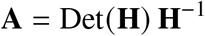

NTs structurally link minima to first-order SP1s, which correspond to transition states in chemical systems [64–66]. Each NT intersects points where:

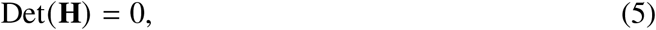

signaling the coalescence of a minimum and saddle point, termed the bond-breaking point (BBP) [56].

In the context of TCR-pMHC interactions, the structural interplay between the hinge angle and the antigen-binding site, as previously discussed, plays a pivotal role in modulating the thermodynamic and kinetic properties of molecular recognition. This interaction is characterized by the non-harmonic coupling between the ligand-binding site separation (*d*) and the conformational dynamics of the protein complex. Consequently, the free energy landscape, which governs both equilibrium states and transition pathways, is primarily dictated by two independent parameters: the protein extension (*L*) and the ligand-binding site distance (*d*). The corresponding geometry (*θ, L, d*) is illustrated in Fig. 3.

**Fig. 3.**
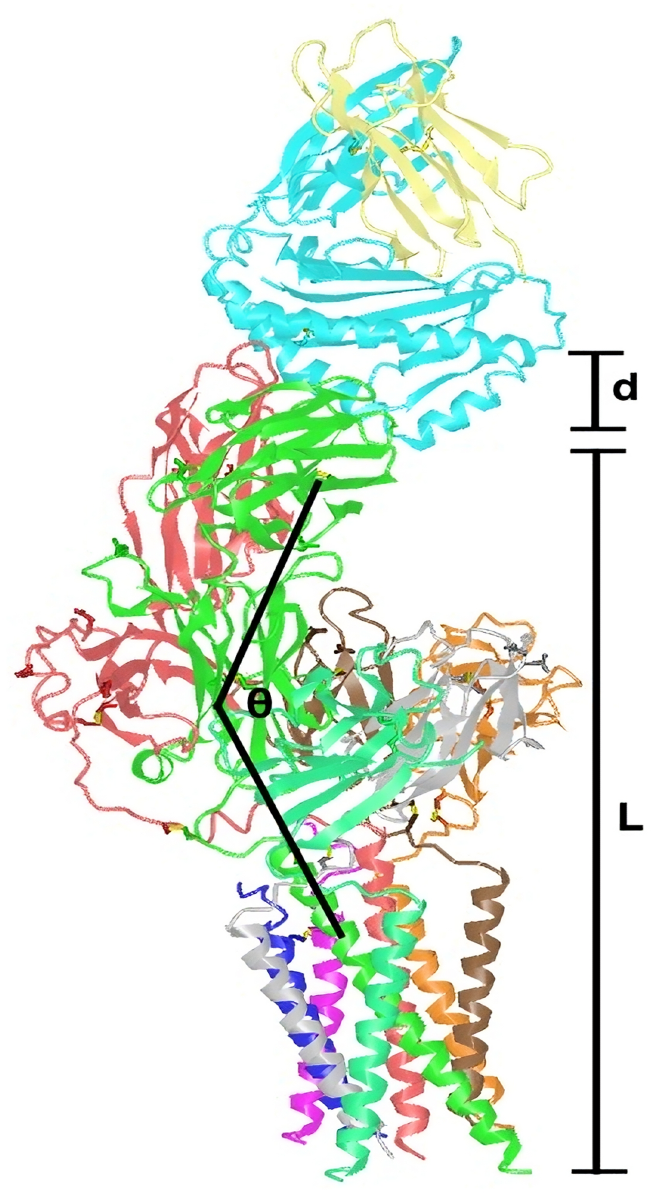
Geometric definitions used in the model. The hinge angle *θ* (black arc) is measured about the TCR ectodomain; the TCR extension *L* is the axial distance from the CD3 transmembrane region to the TCR–pMHC interface; and *d* denotes the ligand-binding site separation along the pulling axis. These coordinates (*L, d, θ*) are those used in the free-energy formulation in Eq. 6.

To enhance analytical tractability, TCR*β* extension, denoted as *L* (*θ*), serves as a surrogate variable for the hinge angle (*θ*), thereby circumventing the need for explicit angular parameterization. This substitution exploits the geometric dependency between *L* and *θ*, which adheres to the functional relationship:

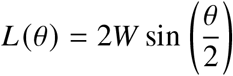

and inversely:

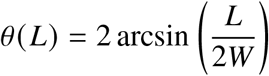

Here, *W* represents the effective length of the TCR complex, estimated at 7 nm [37]. These relationships streamline the representation of the free energy landscape while preserving the essential coupling between conformational dynamics and binding energetics.

The total free energy of the TCR-pMHC system, denoted as *V* _*f*_ (*L, d*), accounts for intrinsic hinge energetics, ligand binding interactions, and external force contributions:

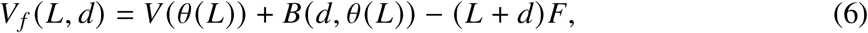

where each term represents a distinct energetic component governing TCR-pMHC interactions [60, 67].

The hinge energy *V* (*θ* (*L*)) describes the system’s preference for a specific hinge angle, modeled using a harmonic potential:

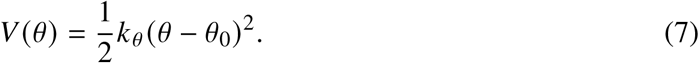

Here, *k*_*θ*_ is the angular stiffness quantifying resistance to deviation from the equilibrium angle *θ*_0_, found to be approximately 0.60*π* radians [38]. The quadratic term penalizes deviations from *θ*_0_, influencing the conformational stability of the TCR [68].

The ligand binding energy *B*(*d, θ* (*L*)) characterizes the interaction between the TCR-pMHC complex, modeled by a Morse potential:

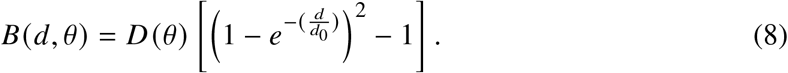

Here, *D* (*θ*) is an angle-dependent binding strength modulating the potential well depth, while *d*_0_ represents the optimal binding distance [69]. The Morse potential captures the non-linear binding behavior, where affinity decreases as *d* increases.

The function *D* (*θ*) reflects allosteric modulation of binding affinity and is given by:

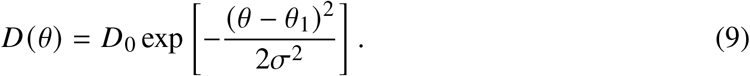

In this formulation, *D*_0_ is the maximum binding strength, *θ*_1_ is the preferred (optimal) hinge angle for binding, and *σ* sets how sharply *D* (*θ*) decays away from *θ*_1_ [70]. The term (*L* + *d*)*F* represents the mechanical work of an applied load *F*, which effectively modulates both the extension *L* and the ligand distance *d*, thereby reshaping the binding landscape [60].

In practice, *F* denotes the effective pico-Newton force transmitted through the TCR–pMHC–CD3 axis (e.g., actin–myosin pulling, membrane tension/curvature, microvillus scanning). Following standard mechanochemistry, for small displacements this transmitted load enters as a linear tilt, −*F* **f**^⊤^**x** with unnormalized **f**^⊤^ = (1, 1) and **x**^⊤^ = (*L, d*) of the PES, capturing how force shifts minima and saddles without specifying its microscopic origin. The constant-load (force-clamp) approximation matches BFP/OT conditions, and the chosen pulling axis reflects microvillus-directed traction. With parameters calibrated to measured pico-Newton ranges, this formulation isolates the effect of force on the potential-energy landscape—specifically, the displacement of minima and saddles—without committing to a particular microscopic origin of the load.

## 3 Mathematical Analysis

We analyze and derive the stationary conditions for bound configurations of the total free-energy equation. Existence and multiplicity are established via a necessary-and-sufficient scalar criterion, and local classification follows from a Hessian (second-derivative) test. We also identify the degeneracy conditions for coalescence of stationary points and provide concise corollaries capturing the relevant sign conditions; full algebraic details are given in the Supplementary Material.

### Theorem 3.1

(Existence and multiplicity of steady states). *Assume W >* 0, *d*_0_ *>* 0, *D*_0_ *>* 0, *σ >* 0, *k*_*θ*_ *>* 0 *and F* ≥ 0. *Let*

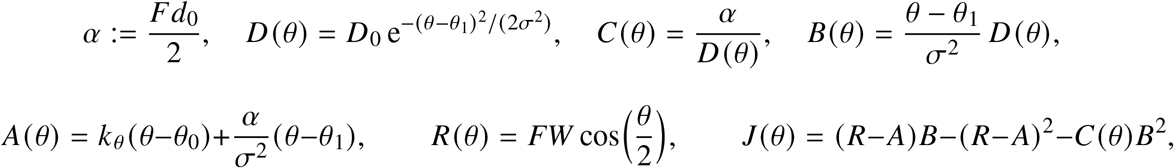

*and define the admissible set* Θ := {*θ* ∈ [0, *π*] : *D* (*θ*) ≥ 2*Fd*_0_} *(nonempty iff D*_0_ ≥ 2*Fd*_0_*). Then:*

i. *If D*_0_ *<* 2*Fd*_0_ *(equivalently, F > D*_0_/(2*d*_0_)*), then* Θ = ∅ *and there are no steady states*.
ii. *If D*_0_ ≥ 2*Fd*_0_, *a pair* (*L*^∗^, *d*^∗^) *is a steady state (a critical point of V* _*f*_ *) if and only if there exists θ*^∗^ ∈ Θ *with J* (*θ*^∗^) = 0. *The corresponding u*^∗^ ∈ (0, 1] *is given by*

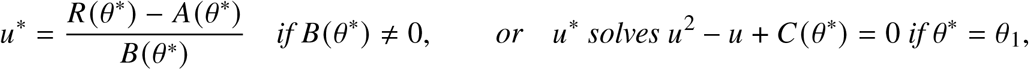

*and then d*^∗^ = −*d*_0_ ln *u*^∗^, *L*^∗^ = 2*W* sin(*θ*^∗^/2). *Moreover, the total number of steady states is at most two*.
iii. *Generically (i*.*e*., *excluding parameter values where a zero of J also satisfies J*^′^(*θ*) = 0*), the number is* 0 *or* 2. *A single steady state occurs only in degenerate situations: either a* 2 *tangency with J* (*θ*) = *J*^′^(*θ*) = 0 *on one side of θ*_1_, *or at the center when*

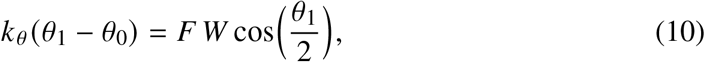

*and D* (*θ*_1_) = 2*Fd*_0_.
iv. *If the center condition* (10) *holds and D* (*θ*_1_) *>* 2*Fd*_0_, *then θ*^∗^ = *θ*_1_ *solves J* = 0 *and both quadratic roots u*_±_ *of u*^2^ − *u* + *C* (*θ*_1_) = 0 *are admissible; thus there are two distinct steady states with the same extension L*^∗^ = 2*W* sin(*θ*_1_/2) *but different separations*.

*Proof (sketch)*. From ∂_*d*_*V* _*f*_ = 0 one obtains *u*(1 − *u*) = *C* (*θ*) with 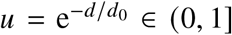, hence *θ* ∈ Θ. Using Φ(*d*) = *u*^2^ −2*u* = −(*C* (*θ*) +*u*), the condition ∂_*L*_*V* _*f*_ = 0 reduces to *A*(*θ*) + *B*(*θ*)*u* = *R*(*θ*). Eliminating *u* yields *J* (*θ*) = 0; conversely, *J* (*θ*^∗^) = 0 with *θ*^∗^ ∈ Θ reconstructs *u*^∗^, then *d*^∗^, *L*^∗^. At *θ*_1_ and at the admissible endpoints *θ*_±_ one finds *J* ≤ 0. Writing *G* = *R* − *A*, one has *J* = −*G*^2^ + *G B* − *C B*^2^ ≤ *J*_max_ := *B*^2^(1/4 − *C*), with *J*_max_ *>* 0 iff *D* (*θ*) *>* 2*Fd*_0_. On each side of *θ*_1_, *J*_max_ is unimodal (product of a single-hump factor 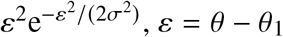, and a strictly decreasing factor 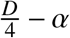), vanishing at side endpoints; thus *J* can cross the axis at most once per side, giving the at most two bound. The one-solution cases follow from tangency *J* = *J*^′^ = 0 on a side or from the center condition together with the threshold *D* (*θ*_1_) = 2*Fd*_0_. If *D* (*θ*_1_) *>* 2*Fd*_0_ under (10), both *u*_±_ are admissible at *θ*_1_, producing two solutions with the same *L*. □

### Proposition 3.2

(Critical force). *Let* 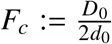. *If F* > *F*_*c*_, *then* Θ = ∅ *and there are no steady states. If* 0 ≤ *F* ≤ *F*_*c*_, *the number of steady states is at most two*.

We now classify local stability of interior stationary points via the Hessian.

### Theorem 3.3

(Local stability at interior stationary points). *Let V* _*f*_ (*L, d*) *be the free energy* (6) *on the interior domain* (*L, d*) ∈ (0, 2*W*) × (0, ∞), *with parameters k*_*θ*_, *D*_0_, *d*_0_ *>* 0. *Assume D* (*θ*) ∈ *C*^2^ *and θ* (*L*) = 2 arcsin(*L*/2*W*) ∈ *C*^2^. *Let* (*L*^∗^, *d*^∗^) *be an interior stationary point:*

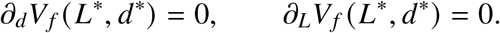

*Set* 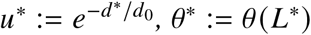, *and evaluate at* (*L*^∗^, *d*^∗^):

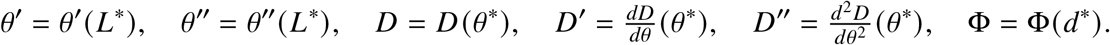

*With variable order* (*L, d*), *the Hessian reads*

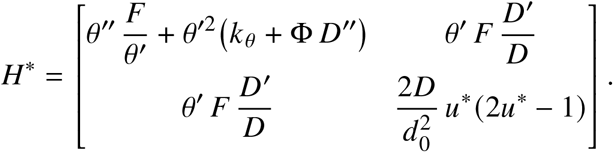

i. ***Necessary curvature in*** *d*.

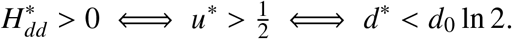 *Hence only the near branch* 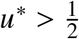 *can be stable along d*.
ii. ***Strict local minimum (classification)***. *For a symmetric* 2 × 2 *Hessian, positive definiteness is equivalent to the positivity of the first leading principal minor and of the determinant (see, e*.*g*., *standard matrix analysis texts [71]):*

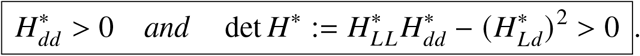 *If* 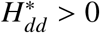 *but* det *H*^∗^ *<* 0 *the point is a saddle; if* 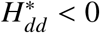 *it is unstable along d*.
iii. ***Zero force*** *F* = 0. *The stationary point is* 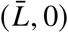 *with* 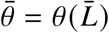 *solving* 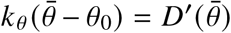; *it is locally stable iff*

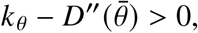

*since* 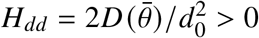 *and H*_*Ld*_ = 0.

Fold remark. *At* 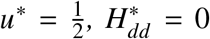 *and* det 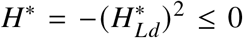 *thus no strict local minimum occurs exactly at the fold*.

### Corollary 3.4

(Sign structure on the *d*-stationary branch). *Along the d-stationary branch, the first-order condition*

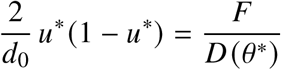

*implies*

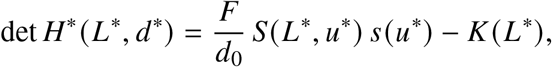

*where*

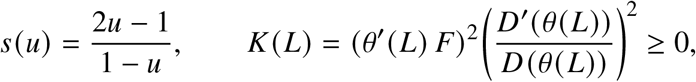

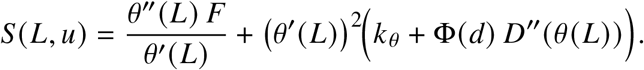

*Consequently:*

- *If S*(*L*^∗^, *u*^∗^) *and s*(*u*^∗^) *have opposite signs, then* det *H*^∗^ *<* 0 *(saddle)*.
- *On the near branch* 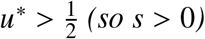, *a strict local minimum requires*

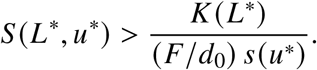

*At the fold* 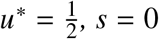 *and* det *H*^∗^ = −*K* ≤ 0, *hence no strict local minimum there*.

Detailed calculations for Theorems 3.1-3.4 are provided in the Supplementary Material.

## 4 Parameter Calculations

**Table 1.** Model parameters for TCR–pMHC angular potential. For all interactions, For all interactions, we assume *W* = 7 nm and *θ*_0_ = 0.60*π* rad based on structural estimates [37, 38]. For reference, *σ* in radians equals 0.222 (agonist), 0.254 (weak), and 0.482 (antagonist).

| Nature | $k_\theta$<br>(pN·nm) | $D_0$<br>(pN·nm) | $d_0$<br>(nm) | $\sigma/\pi$ | $\theta_1/\pi$ |
| --- | --- | --- | --- | --- | --- |
| Agonist | 76.3 | 51.41 | 0.185 | 0.0707 | 0.83 |
| Weak Agonist | 80.9 | 40.30 | 0.151 | 0.08095 | 0.80 |
| Antagonist | 85.3 | 31.47 | 0.125 | 0.15335 | 0.72 |

We quantify binding strength as a positive well depth *D*_0_ (per molecule). For each peptide–TCR pair, *D*_0_ is obtained from its 3D dissociation constant *K*_*D*_ via the standard-state relation

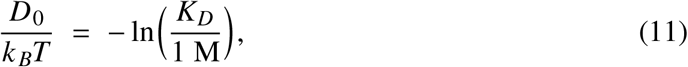

where ln is the natural logarithm, 1 M is the standard state, and *T* is the assay temperature (we use *T* = 310*K*). When a molar free energy is reported instead, we use the equivalent one-molecule definition *D*_0_ = −Δ*G*^°^/*N*_*A*_. We report *D*_0_ both as the dimensionless ratio *D*_0_/*k* _*B*_*T* and in pN·nm using 1 pN · nm = 10^−21^ J; at *T* = 310*K, k* _*B*_*T* = 4.28 pN·nm. At *T* = 310 K, the corresponding *D*_0_/*k* _*B*_*T* values are 12.0 (agonist), 9.4 (weak), and 7.35 (antagonist). This sign convention matches our energy landscape: larger *D*_0_ corresponds to a deeper potential well and stronger binding.

In practice, *K*_*D*_ (and therefore *D*_0_) is ligand-specific even within the same functional class. Accordingly, we compute *D*_0_ per peptide and use class labels (agonist, weak/partial agonist, antagonist) only to summarize the resulting distributions (median and interquartile range). To avoid cross-study mixing of energy scales, our main parameterization relies exclusively on 3D (solution) affinities from the same experimental sources for each dataset.

Consistent with prior measurements of TCR–pMHC affinities [72–74], the per-ligand *D*_0_ values in our 3D compilations fall into characteristic ranges: agonists typically ∼ 48–55 pN·nm, weak/partial agonists ∼ 38–41 pN·nm, and antagonists ∼ 31–35 pN·nm. For concreteness we include representative ligands across this spectrum—gp33-wt (agonist) [74], Y4A (weak agonist) [72], and G4Y (antagonist) [74]. Here, *D*_0_ represents an effective one-coordinate well depth along the ligand separation axis that reproduces the measured three-dimensional free energy Δ*G*^°^ after integrating out other degrees of freedom, thereby ensuring thermodynamic–mechanical consistency in our parameterization. Full per-ligand conversions (Δ*G* → *D*_0_) and class summaries for each dataset are provided in the Supplementary Information.

The angular stiffness (*k*_*θ*_) for each ligand class was estimated by fitting a unified nonlinear model to experimentally measured bond lifetimes as a function of applied force [53], together with angular displacement parameters derived from structural analyses of TCR–pMHC–CD3 complexes. The model links the mean lifetime *τ*(*F*) to both angular deformation and force-dependent stabilization, allowing the distinct mechanical signatures of agonist, weak agonist, and antagonist interactions to emerge directly from the data. Parameter estimation was performed by nonlinear least-squares optimization in Matlab (R2024b) with weak biophysical priors, and parameter uncertainties were quantified via residual bootstrapping.

The resulting stiffness estimates were *k*_*θ*_ for the agonist = 76.3 [72.8, 79.7] pN·nm, for the weak agonist = 80.9 [73.8, 88.0] pN·nm, and for the antagonist = 85.3 [61.6, 109.1] pN·nm, consistent with progressively stiffer angular potentials from agonist to antagonist complexes. For each applied force, the bound and transition states were located on the two-dimensional potential surface *V* _*f*_ (*L, d*) to compute the energy barrier Δ*G*^‡^ (*F*), and bond lifetimes were fitted using a Kramers-type relation with a single attempt frequency.

We model the angle dependence of the binding depth as

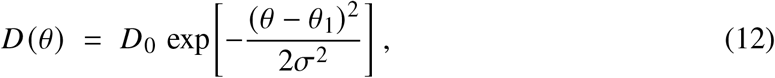

where *σ* sets the *intrinsic angular tolerance* of the binding interaction (smaller *σ* = sharper selectivity near *θ*_1_; larger *σ* = broader tolerance). Because the observed bound spread also reflects hinge elasticity (near the minimum, the effective curvature is *κ*_eff_ = *k*_*θ*_ + *D*_0_/*σ*^2^), we do *not* equate *σ* to a measured bound FWHM except for a single anchor point.

The angular width parameter *σ* defines the effective angular tolerance of the TCR–pMHC interface and directly modulates the breadth of the binding potential *D* (*θ*). To parameterize this quantity consistently across ligands, we first relate experimentally observed angular spreads to model parameters and then link them to binding thermodynamics.

For each dataset, we define a reference width *σ*_0_ from a well-characterized agonist by converting a reported full width at half maximum (FWHM) to a Gaussian standard deviation,

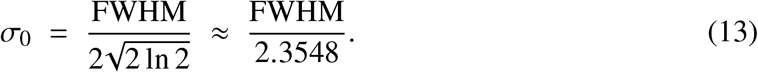

Using a commonly reported bound width of 30^°^ for the TCR–CD3–pMHC complex [38] gives *σ*_0_ ≃ 0.222 rad.

To capture ligand-to-ligand variability within the same study, we scale *σ* monotonically with the molar binding free energy (Δ*G*) using a Boltzmann-like relation,

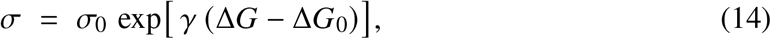

where Δ*G*_0_ is the free energy of the reference agonist and *γ* has units (kcal mol^−1^)^−1^. This phenomenological form follows the statistical–mechanical relation between energetic stability and configurational variance [53, 75–77]. The parameter *γ* is determined within each dataset by selecting a target angular tolerance for a representative antagonist, ensuring that *σ* remains physically realistic and monotonic across all ligands. The Δ*G* values are taken directly from the experimental sources reporting the corresponding peptide–MHC interactions [72–74, 78]. Full calibration details (reference peptide, Δ*G*_0_, target width, fitted *γ*, and per-ligand *σ* values) are provided in the Supplementary Information.

The angle *θ*_1_ denotes the reference or central angle at which the binding strength function *D* (*θ*) reaches its maximum. It corresponds to the optimal angular configuration of the receptor–ligand complex, representing the orientation that yields the strongest interaction. This central angle thus defines the most favorable geometrical alignment for productive binding. Deviations from *θ*_1_ lead to a reduction in binding strength, with the rate of decay modulated by the spread parameter *σ*, which governs the system’s angular tolerance. Using the extension–angle relation *θ*_1_ = 2 arcsin(*L*_bound_/2*W*) with *W* = 7 nm, the resulting angular preferences were set to *θ*_1_/*π* = 0.83 (Agonist), 0.80 (Weak Agonist), and 0.72 (Antagonist), reflecting progressively more closed hinge conformations across the affinity series. These choices are consistent with cryo-EM comparisons indicating an opening toward ∼ 150^°^ upon pMHC engagement [38].

Experimental work shows that, upon engagement with pMHC, the TCR–CD3 complex adopts a more extended, open conformation, with peak interactions typically near 150^°^ [38]. To compare directly with these observations, we measured the TCR*β* tilt angle in unbound and pMHC-bound states using two high-resolution cryo-EM structures: the unbound TCR from Dong *et al*. [40] and the pMHC-bound complex from Saotome *et al*. [45]. For consistency, the hinge angle was defined by the same three residues in both structures—VAL_3_, GLY_164_, and ARG_289_—taken as the angle subtended by VAL_3_–GLY_164_–ARG_289_. Using UCSF Chimera [79], we obtained a tilt of 104.99^°^ in the unbound state and 147.56^°^ in the pMHC-bound state, indicating a substantial opening upon antigen engagement (Appendix Fig. B1). This shift is consistent with Eerden *et al*. [38], who report a more extended TCR configuration near 150^°^. All structures were retrieved from the National Library of Medicine Protein Data Bank [80]. Moreover, the magnitude of this conformational change appears to correlate with antigen potency: strong agonists tend to induce larger extensions than weaker ligands, in line with enhanced binding specificity and mechanical stabilization.

The parameter *W* represents the length of the TCR-CD3 complex, estimated at 7 nm. It defines the relationship between the angle of the hinge *θ* and the extension *L* (*θ*), allowing *L* to capture conformational changes in the complex. These conformational changes are essential in modulating the binding dynamics and energy landscapes of the system. The parameter *d*_0_, representing the interaction range between TCR and pMHC, is adopted following the characterization provided in [81].

## 5 Results

### 5.1 From Catch to Slip: Force-Programmed Barriers in TCR–pMHC

We examine the impact of external forces on TCR bound with pMHC that has an agonist, weak agonist, and antagonist peptide. An agonist binds to a receptor and activates it, triggering a biological response [82, 83]. A weak agonist, as described by [84], also binds but elicits only a partial response, even at full receptor occupancy. An antagonist binds without activation, thus blocking the effects of the agonist [85]. This competitive inhibition was then quantified through dose-response analysis [86].

We examined the influence of an externally applied force oriented along the (1, 1) direction across all ligand classes, focusing on the distinct energetic responses arising from different molecular interactions. Based on the parameter values presented in the previous section, the agonist case displays a non-monotonic dependence of the energy barrier on the applied force. As the force increases, the barrier height rises up to 9 pN, where it reaches a maximum, before decreasing at higher forces. This biphasic pattern is characteristic of a catch–slip bond, in which moderate tensile forces enhance bond stability through barrier strengthening, whereas larger forces promote bond dissociation. This observation is consistent with experimental reports of TCR–pMHC interactions, where force-enhanced stability occurs within an intermediate force range before rupture becomes dominant at high force [10]. A similar, though less pronounced, response is observed in the weak agonist case. Here, the energy barrier increases initially, peaking around 9 pN, followed by a gradual decline at higher forces. This indicates a weaker catch–slip phenotype, consistent with the reduced mechanical reinforcement reported for altered peptide ligands [23]. In contrast, the antagonist case exhibits a monotonic decrease of the energy barrier with increasing force. No intermediate stabilization is observed; instead, the barrier continuously diminishes as the applied force grows. This force-weakening profile reflects a pure slip-bond behavior, where external force directly facilitates bond dissociation. Such a response mirrors experimental force-spectroscopy observations for non-stimulatory TCR–pMHC interactions, which lack the force-induced stabilization characteristic of agonist complexes [53]. The detailed numerical values corresponding to these trends are provided in the Tables in Appendix (A1,A2,A3).

The bond dissociation threshold varies across interaction types, occurring at approximately 116 pN for the agonist, 105 pN for the weak agonist, and 75 pN for the antagonist. At these critical force levels, the transition state (TS) and the energy minimum (R) merge at the BBP, marking the irreversible dissociation of the system. This phenomenon underscores the intricate interplay between force magnitude and molecular stability, providing key insights into the mechanistic principles governing catch-slip bond behavior.

The PES diagrams (6) illustrate how the TCR extension (*L*) and the distance to the ligand (*d*) affect the stability of the TCR-pMHC bond under varying force conditions. Each contour plot represents the equipotential lines of the potential energy function in the (*L, d*) coordinate space, where the two length coordinates (*L* and *d*, in nm) describe the movement of the ligand.

The Reactant (R) corresponds to the global minimum of the PES, representing the bound state of the ligand-receptor interaction. The TS is a SP1 of index one, signifying the energy peak that must be overcome for bond dissociation. The BBP marks the moment when an applied force causes the former minimum (R) and the SP1 to coincide, resulting in bond rupture. The Det(**H**) = 0 lines, shown in green, indicate critical points where the determinant of the Hessian matrix vanishes, signifying structural transitions in the energy landscape that impact molecular stability and interactions.

The bold curve is the NT. On it, the minimum and SP1 move. After the minimum, because of the direction, (1, 1), the NT ascends strongly into the PES mountains for higher *L* values. This causes also an increasing barrier height, thus a catch bond behavior. We plotted the PES for different force values, considering the cases of an agonist, weak agonist, and antagonist.

For the agonist in the forceless limit (*F* = 0), the PES is represented by thin equipotential lines, with R as the global minimum, indicating the bound state. The TS, corresponding to the dissociation limit, lies far beyond the displayed scale. The thick Newton trajectory (NT) along the (1,1) direction represents a slip-catch-slip bond transition. The Det(**H**) = 0 lines, shown in green, highlight critical points in the PES where the determinant of the Hessian matrix vanishes, signaling structural transitions in the energy landscape. For the case of *F* = 10 pN, we observe a single minimum (R) and a single SP. Notably, the solid NT has a TP, which may serve as an indicator of a catch bond mechanism, suggesting force-enhanced binding stability. Finally, at a high force of *F* = 116 pN, the former TS and the former minimum (R) merge to form a shoulder, indicating the disappearance of the energy barrier.

For the weak agonist, in a similar way for the case of *F* = 10 pN, we observe a single minimum (R) and a single saddle point (SP). Notably, the solid NT has a TP, which may serve as an indicator of a catch bond mechanism, suggesting force-enhanced binding stability. At a high force of *F* = 105 pN, the former TS and the former minimum (R) merge to form a shoulder, indicating the disappearance of the energy barrier.

In conclusion, forces applied along the (1, 1) direction reveal distinct, peptide-class–specific mechanical responses. Agonists exhibit a characteristic catch–slip profile, in which the energy barrier increases up to approximately 9 pN before declining at higher loads. Weak agonists show a similar but attenuated peak near 9 pN, whereas antagonists behave as pure slip bonds, displaying a monotonic barrier decay. Bond-rupture events occur when the bound minimum and transition state merge, at roughly 116 pN for agonists, 105 pN for weak agonists, and 75 pN for antagonists. Analysis of the PES and Newton trajectories—organized by lines where Det(**H**) = 0 —links these outcomes to the underlying force-reshaped energy landscapes in (*L, d*) space. Moderate mechanical loads can transiently stabilize agonist binding, whereas larger forces universally promote bond dissociation across all classes.

### 5.2 Force-Dependent Bond Lifetimes

The mean bond lifetime, particularly in the context of adhesion, unbinding, and bond stability, is influenced by an energy barrier that must be overcome for bond dissociation. Theoretical models such as the Arrhenius kinetics and Kramers’ rate theory offer a framework to quantify bond dissociation under thermal fluctuations and external forces [87, 88]. In the overdamped (Smoluchowski) regime, the lifetime of the bond, *τ*, is commonly estimated using the Arrhenius-type equation:

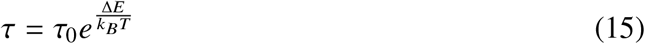

where Δ*E* represents the energy barrier, *k* _*B*_ is the Boltzmann constant, *T* is the absolute temperature, and *τ*_0_ is a prefactor that accounts for the attempt frequency of escape from the potential well [89].

This energy barrier Δ*E* can also be represented by Δ*E* = *V* _*f*_ (**x**_*s*_) − *V* _*f*_ (**x**_*m*_) where the vectors **x**_*m*_ = (*L*_*m*_, *d*_*m*_) and **x**_*s*_ = (*L*_*s*_, *d*_*s*_) denote, respectively, the positions in configuration space of the local minimum (the bound state) and saddle point (the transition state) of *V* _*f*_. At human body temperature, the thermal energy scale is given by *k* _*B*_*T* ≈ 4.28 pN · nm which provides a natural unit for comparing energy barriers in molecular interactions.

The pre factor *τ*_0_ for the escape rate in Langer’s [90] metastable decay theory is given by:

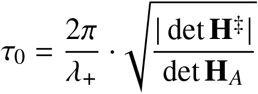

Here, *λ*_+_ is given by *λ*_+_ = −*β* 𝒟 *h*_*uu*_, where *β* is the inverse thermal energy, defined as 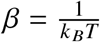, with *k* _*B*_ being the Boltzmann constant and *T* the temperature in Kelvin. The parameter 𝒟 denotes the (isotropic) diffusion coefficient (units nm^2^/s; related to friction by the Einstein relation 𝒟 = *k* _*B*_*T* /*ζ*), which accounts for frictional effects in Kramers’ theory. The term det **H**_*A*_ denotes the determinant of the Hessian matrix at the local minimum, while det **H**^‡^ represents the determinant of the Hessian matrix at the saddle point (the absolute value is taken because an index-1 saddle has det **H**^‡^ *<* 0). Additionally, 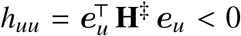 is the unstable curvature (negative eigenvalue) of the saddle Hessian along the unit unstable direction ***e***_*u*_.

The bond lifetime (*τ*) as a function of applied force was analyzed for three ligand types: agonist, weak agonist, and antagonist. Each ligand exhibited a distinct bond lifetime in response to increasing force (see Figure 7). Analysis of the computed lifetimes (*τ*) as a function of applied force reveals distinct mechanochemical behaviors across the three peptide classes. In the agonist case, bond lifetime increases from 0.83 s at 1 pN to a maximum of approximately 1.22 s at 8–9 pN, beyond which it gradually declines to 0.24 s by 30 pN. This biphasic trend indicates a catch–slip transition, where moderate tensile forces stabilize the complex through an increased energy barrier, while higher forces promote bond rupture. For the weak agonist, the lifetime initially decreases from 0.16 s at 1 pN to 0.15 s at 3 pN, followed by a shallow rise to about 0.20 s near 8–9 pN before decreasing again to 0.04 s at 30 pN. This subtle peak suggests a weak or transient catch–bond behavior, with limited mechanical reinforcement compared with the full agonist. By contrast, the antagonist exhibits a strictly monotonic decay in lifetime with increasing force, characteristic of a pure slip bond, where external load directly facilitates dissociation rather than stabilization.

**Fig. 4.**
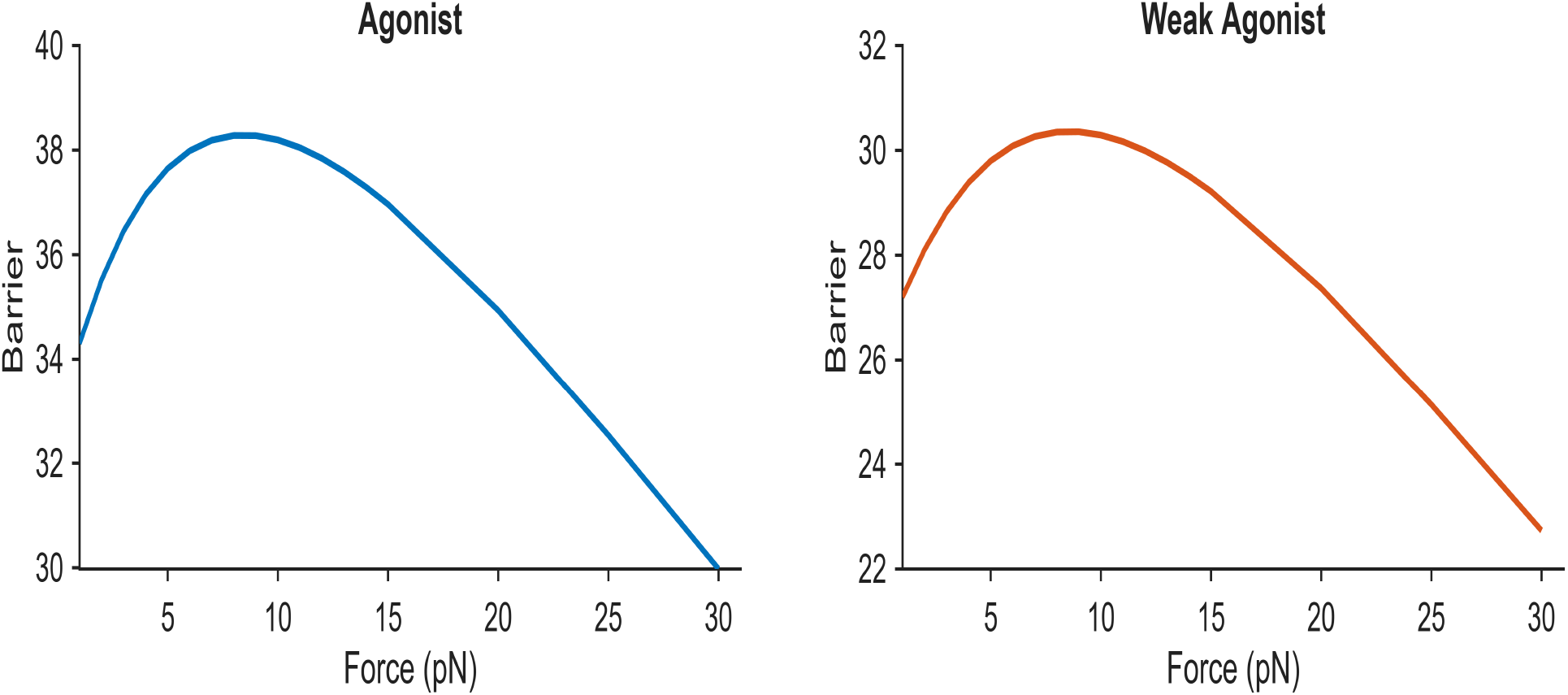
Plot for Barrier vs Force in case of agoinst and weak agonist.

**Fig. 5.**
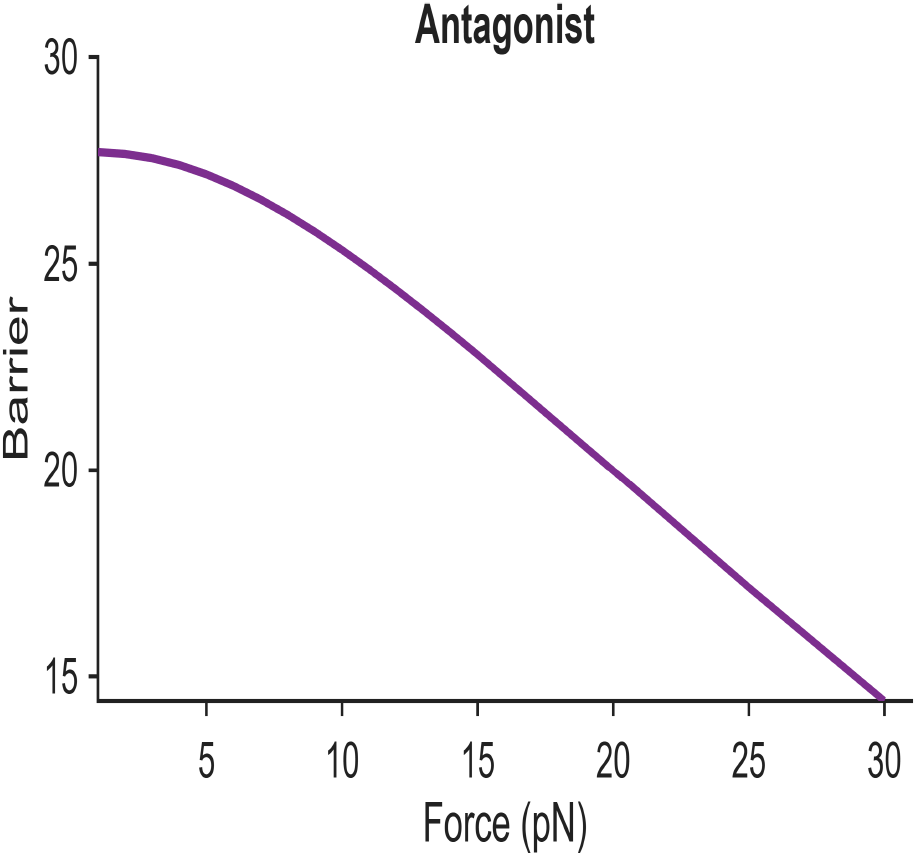
Plot for Barrier vs Force in case of antagonist.

**Fig. 6.**
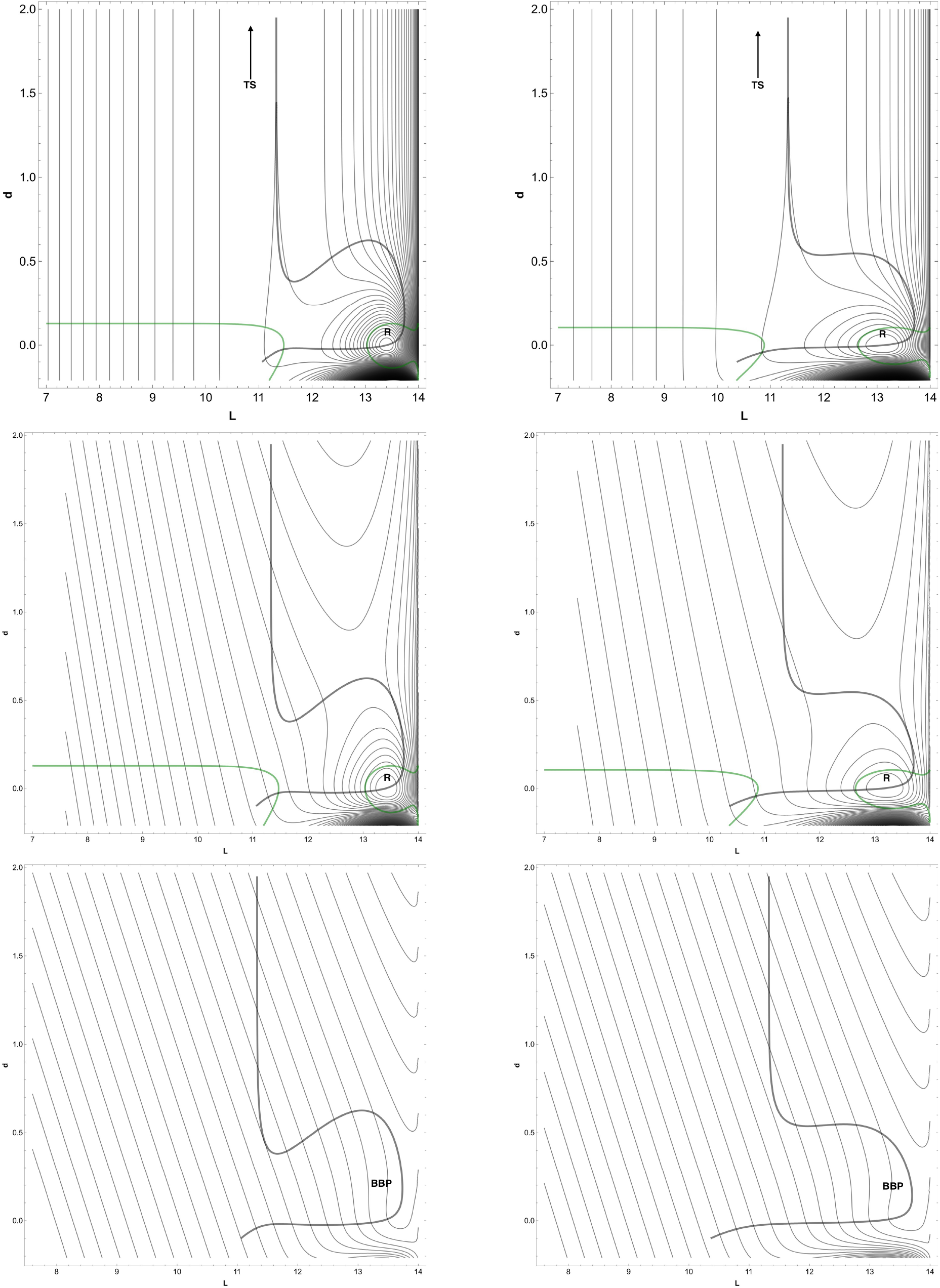
2D potential energy surfaces (PES) for TCR–pMHC interactions under different forces. The left column corresponds to the agonist, and the right column to the weak agonist. From top to bottom: *F* = 0, *F* = 10, and high-force conditions (*F* = 116 for agonist and *F* = 105 for weak agonist). Equipotential contours represent energy landscapes under each condition.

**Fig. 7.**
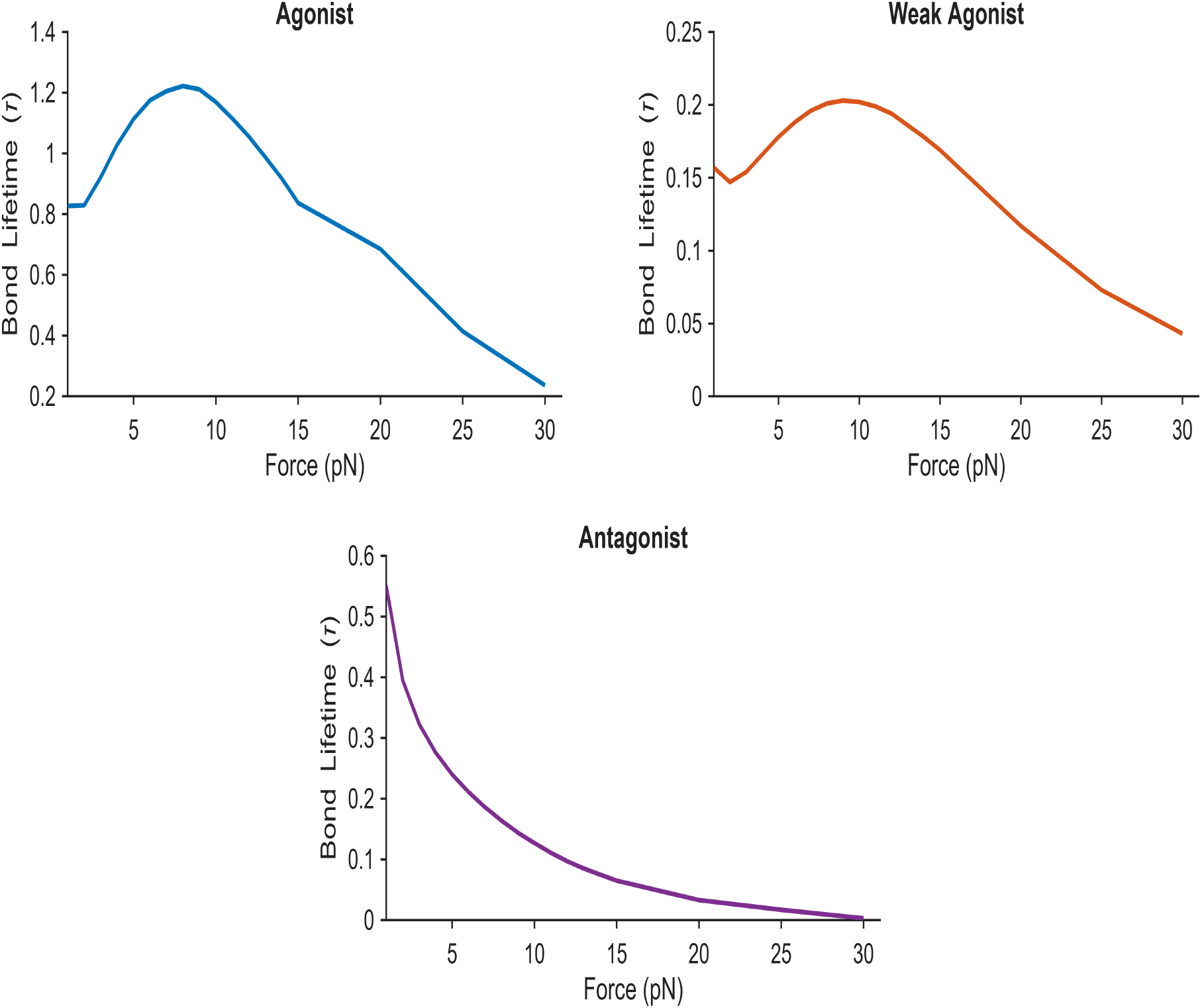
Bond lifetime (*τ*) as a function of force (*F*) for agonist (top left), weak agonist (top right), and antagonist (bottom). The agonist shows catch-slip behavior with with lifetime peaking around 8–9 pN. The weak agonist exhibits a mild or transient stabilization, indicating limited mechanical reinforcement, where as the antagonist exhibits a monotonic decrease with pure slip bond behavior.

Overall, only agonists display a pronounced catch–slip profile, consistent with their ability to enhance bond stability under physiological force ranges and thereby promote TCR activation. Weak agonists show marginal or inconsistent stabilization, while antagonists destabilize under load, reflecting their inhibitory role in T-cell signaling.

## 6 Conclusions

Within a mechanochemical framework, we apply Newton-trajectory analysis to our loaded PES to track minima–saddle motion and identify bond-breaking points, and then use Arrhenius–Kramers theory to predict the associated bond lifetimes. By applying Newton-trajectory theory to the loaded potential-energy surface *V* _*f*_ (cf. (6)), we systematically follow how minima and saddles migrate under load, identify the precise bond-breaking point (where they coalesce), establish an existence–stability threshold for steady states, and link force-driven barrier reshaping to experimentally observed bond-lifetime patterns—catch–slip for agonists and weak agonists versus slip for antagonists—across peptide classes. We identify a sharp threshold *F*_*c*_ = *D*_0_/(2*d*_0_) (see 3.2): for *F > F*_*c*_ no steady states exist, while for 0 ≤ *F* ≤ *F*_*c*_ there are at most two (typically 0 or 2). Generically, solutions appear as a paired minimum–saddle; a single solution arises only in degenerate parameter choices (3.1). When steady states exist, they are found by solving *J* (*θ*) = 0 on the admissible set Θ, then reconstructing *u, d*, and *L* (3.1). Local stability is possible only on the “near” branch *u >* 1/2 with det *H >* 0; at the fold *u* = 1/2 the minimum and saddle coalesce (BBP) and stability is lost(3.3,3.4).

We calibrated model parameters directly from data in a manner consistent with 4. Binding strength *D*_0_ was derived from reported *K*_*D*_ values via the standard-state relation *D*_0_ = −*k*_*B*_*T* ln *K*_*D*_/1 M ; the angular width *σ* came from the FWHM of hinge-angle distributions (with any ligand-to-ligand scaling by Δ*G*); and the angular stiffness *k*_*θ*_ was estimated by fitting force-dependent bond lifetimes using an Arrhenius–Kramers form. The interaction range *d*_0_ used class-specific literature estimates, and geometry was fixed by the cryo-EM length scale *W* with the standard *θ* ↔ *L* map. These calibrated inputs populate the Gaussian angular binding term *D* (*θ*) and the Morse distance term *B*(*d, θ*) used in the energy landscape. With these inputs, we establish class-specific mechanical phenotypes: agonist and weak-agonist peptides exhibit catch–slip behavior in their energy barriers (barriers increase with moderate load then decrease), while antagonists display slip behavior with steadily falling barriers; correspondingly, bond lifetimes peak at intermediate load for agonists (with a smaller peak for weak agonists) and decline continuously for antagonists.

Our approach combines a minimal, data-anchored energy landscape (calibrated from *K*_*D*_, cryo-EM angles, and literature scales) with Newton-trajectory analysis to prove an existence stability threshold and pinpoint the bond-breaking point. Unlike simulation-heavy pipelines, it yields closed-form, interpretable criteria (e.g., *F*_*c*_ = *D*_0_/(2*d*_0_); stability only for *u >* 1/2) and directly links barrier reshaping to bond lifetimes. This delivers class-specific, testable predictions (catch–slip for agonists/weak agonists; slip for antagonists) in a compact, mechanistically transparent framework.

Our parameter choices (e.g., *D*_0_ from *K*_*D*_, class-specific *d*_0_, and *σ* from FWHM with optional empirical scaling in the Supplementary Information) provide a practical first calibration, though they naturally carry some uncertainty for barrier and lifetime estimates. The model is deliberately minimal—Gaussian angular approximation and constant loading—to keep the physics transparent, which means effects such as membrane mechanics, viscoelastic response, and peptide-specific shifts in *θ*_1_ are not yet included. Partnering with experimentalists to obtain peptide- and system-specific measurements for these parameters would refine calibration and increase predictive power. With such data, the framework can be extended to circular angle statistics, time-dependent loading, and membrane-coupled effects to better mirror cellular conditions. In sum, our framework offers a clear, testable mechanochemical basis for TCR recognition—linking force, barriers, and lifetimes—and a practical roadmap for refining predictions with targeted measurements to better anticipate T-cell signaling under load.

## Appendix A. Summary of stationary points and barrier heights for agonist, weak agonist, and antagonist ligands

This appendix provides the numerical results corresponding to the stationary points and transition-state energies of the effective potential *V*_eff_ governing the TCR–pMHC interaction. For each ligand class—agonist, weak agonist, and antagonist—the equilibrium minima and saddle points were computed along the normal–tangential (NT) reaction coordinate 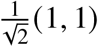. The tables below list the values of the extension *L*, ligand separation *d*, and the corresponding potential energies (in pN·nm), from which the barrier heights were derived. These data form the quantitative basis for the force-dependent bond lifetimes and transition analyses discussed in the main text.

### Agonist

**Table A1.** Stationary points of the effective potential *V*_eff_ for the TCR–pMHC interaction, obtained along the NT direction 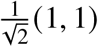. Energies are given in pN·nm.

| $F$ | Min ($L, d$) [nm] | $V_{\text{Min}}$ [pN] | SP ($L, d$) [nm] | $V_{\text{SP}}$ [pN] | Barrier |
| --- | --- | --- | --- | --- | --- |
| 1 | (13.4101, 0.0003) | −46.2752 | (11.6634, 0.3804) | −11.9705 | 34.3047 |
| 2 | (13.4139, 0.0007) | −59.6878 | (11.9297, 0.4167) | −24.1668 | 35.5210 |
| 3 | (13.4177, 0.0010) | −73.1045 | (12.1439, 0.4676) | −36.6497 | 36.4548 |
| 4 | (13.4214, 0.0014) | −86.5253 | (12.3204, 0.5114) | −49.3748 | 37.1505 |
| 5 | (13.4251, 0.0017) | −99.9501 | (12.4680, 0.5459) | −62.3005 | 37.6496 |
| 6 | (13.4287, 0.0021) | −113.3789 | (12.5930, 0.5721) | −75.3924 | 37.9865 |
| 7 | (13.4323, 0.0024) | −126.8116 | (12.6998, 0.5915) | −88.6225 | 38.1891 |
| 8 | (13.4358, 0.0028) | −140.2483 | (12.7919, 0.6054) | −101.9684 | 38.2799 |
| 9 | (13.4393, 0.0031) | −153.6887 | (12.8720, 0.6150) | −115.4082 | 38.2805 |
| 10 | (13.4427, 0.0035) | −167.1330 | (12.9421, 0.6210) | −128.9378 | 38.1952 |
| 11 | (13.4461, 0.0038) | −180.5811 | (13.0039, 0.6243) | −142.5343 | 38.0468 |
| 12 | (13.4495, 0.0042) | −194.0328 | (13.0586, 0.6255) | −156.1912 | 37.8416 |
| 13 | (13.4528, 0.0045) | −207.4883 | (13.1075, 0.6249) | −169.9000 | 37.5883 |
| 14 | (13.4560, 0.0049) | −220.9474 | (13.1512, 0.6229) | −183.6537 | 37.2937 |
| 15 | (13.4593, 0.0052) | −234.4101 | (13.1906, 0.6199) | −197.4464 | 36.9637 |
| 20 | (13.4750, 0.0071) | −301.7768 | (13.3396, 0.5947) | −266.8417 | 34.9351 |
| 25 | (13.4899, 0.0090) | −369.2294 | (13.4373, 0.5624) | −336.6935 | 32.5359 |
| 30 | (13.5042, 0.0109) | −436.7647 | (13.5056, 0.5291) | −406.7888 | 29.9759 |

### Weak Agonist

**Table A2.** Stationary Points of the potential energy surface *V*_eff_ for the TCR–pMHC interaction by Eq. 1, along the NT direction 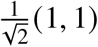. Energies are in pN·nm.

| $F$ | Min ($L, d$) [nm] | $V_{\text{Min}}$ [pN] | SP ($L, d$) [nm] | $V_{\text{SP}}$ [pN] | Barrier |
| --- | --- | --- | --- | --- | --- |
| 1 | (13.1523, 0.0003) | -39.3487 | (11.5877, 0.5744) | -12.1439 | 27.2048 |
| 2 | (13.1613, 0.0006) | -52.5060 | (11.8089, 0.5407) | -24.3981 | 28.1079 |
| 3 | (13.1699, 0.0009) | -65.6723 | (11.9974, 0.5368) | -36.8414 | 28.8309 |
| 4 | (13.1784, 0.0012) | -78.8476 | (12.1592, 0.5395) | -49.4596 | 29.3880 |
| 5 | (13.1867, 0.0015) | -92.0315 | (12.2991, 0.5431) | -62.2318 | 29.7997 |
| 6 | (13.1948, 0.0018) | -105.2239 | (12.4210, 0.5459) | -75.1378 | 30.0861 |
| 7 | (13.2027, 0.0021) | -118.4245 | (12.5278, 0.5473) | -88.1601 | 30.2644 |
| 8 | (13.2105, 0.0024) | -131.6334 | (12.6220, 0.5473) | -101.2834 | 30.3500 |
| 9 | (13.2181, 0.0027) | -144.8502 | (12.7056, 0.5460) | -114.4948 | 30.3554 |
| 10 | (13.2255, 0.0029) | -158.0748 | (12.7801, 0.5435) | -127.7832 | 30.2916 |
| 11 | (13.2328, 0.0032) | -171.3071 | (12.8468, 0.5402) | -141.1391 | 30.1680 |
| 12 | (13.2400, 0.0035) | -184.5469 | (12.9068, 0.5360) | -154.5546 | 29.9923 |
| 13 | (13.2471, 0.0038) | -197.7941 | (12.9610, 0.5312) | -168.0226 | 29.7715 |
| 14 | (13.2540, 0.0042) | -211.0486 | (13.0101, 0.5259) | -181.5371 | 29.5115 |
| 15 | (13.2608, 0.0045) | -224.3104 | (13.0549, 0.5201) | -195.0930 | 29.2174 |
| 20 | (13.2933, 0.0060) | -290.7229 | (13.2283, 0.4881) | -263.3533 | 27.3696 |
| 25 | (13.3237, 0.0076) | -357.3004 | (13.3458, 0.4543) | -332.1620 | 25.1384 |
| 30 | (13.3523, 0.0093) | -424.0334 | (13.4296, 0.4215) | -401.3006 | 22.7328 |

### Antagonist

**Table A3.** Stationary points on the potential energy surface *V*_eff_ of the TCR–pMHC complex (Eq. 1), along the normal transition (NT) direction 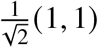. Energies are given in pN·nm.

| $F$ | Min ($L, d$) [nm] | $V_{\text{Min}}$ [pN] | SP ($L, d$) [nm] | $V_{\text{SP}}$ [pN] | Barrier |
| --- | --- | --- | --- | --- | --- |
| 1 | (12.2427, 0.0003) | -39.9972 | (11.5232, 0.7483) | -12.2956 | 27.7016 |
| 2 | (12.2963, 0.0005) | -52.2673 | (11.6993, 0.6688) | -24.6122 | 27.6551 |
| 3 | (12.3468, 0.0008) | -64.5897 | (11.8573, 0.6238) | -37.0367 | 27.5530 |
| 4 | (12.3948, 0.0010) | -76.9616 | (11.9993, 0.5922) | -49.5735 | 27.3881 |
| 5 | (12.4403, 0.0013) | -89.3805 | (12.1274, 0.5677) | -62.2174 | 27.1631 |
| 6 | (12.4838, 0.0015) | -101.8441 | (12.2431, 0.5474) | -74.9609 | 26.8832 |
| 7 | (12.5253, 0.0018) | -114.3505 | (12.3480, 0.5299) | -87.7958 | 26.5547 |
| 8 | (12.5650, 0.0020) | -126.8976 | (12.4434, 0.5143) | -100.7142 | 26.1834 |
| 9 | (12.6031, 0.0023) | -139.4840 | (12.5303, 0.5002) | -113.7088 | 25.7752 |
| 10 | (12.6397, 0.0026) | -152.1079 | (12.6097, 0.4872) | -126.7730 | 25.3349 |
| 11 | (12.6749, 0.0028) | -164.7680 | (12.6824, 0.4751) | -139.9007 | 24.8673 |
| 12 | (12.7088, 0.0031) | -177.4629 | (12.7491, 0.4638) | -153.0863 | 24.3766 |
| 13 | (12.7415, 0.0034) | -190.1914 | (12.8105, 0.4531) | -166.3249 | 23.8665 |
| 14 | (12.7731, 0.0036) | -202.9523 | (12.8671, 0.4428) | -179.6119 | 23.3404 |
| 15 | (12.8037, 0.0039) | -215.7446 | (12.9194, 0.4331) | -192.9434 | 22.8012 |
| 20 | (12.9426, 0.0054) | -280.1424 | (13.1295, 0.3897) | -260.1512 | 19.9912 |
| 25 | (13.0623, 0.0071) | -345.1928 | (13.2786, 0.3528) | -328.0457 | 17.1471 |
| 30 | (13.1668, 0.0089) | -410.8113 | (13.3883, 0.3205) | -396.4078 | 14.4035 |

## Appendix B.

### Supplementary Figures

**Fig. B1.**
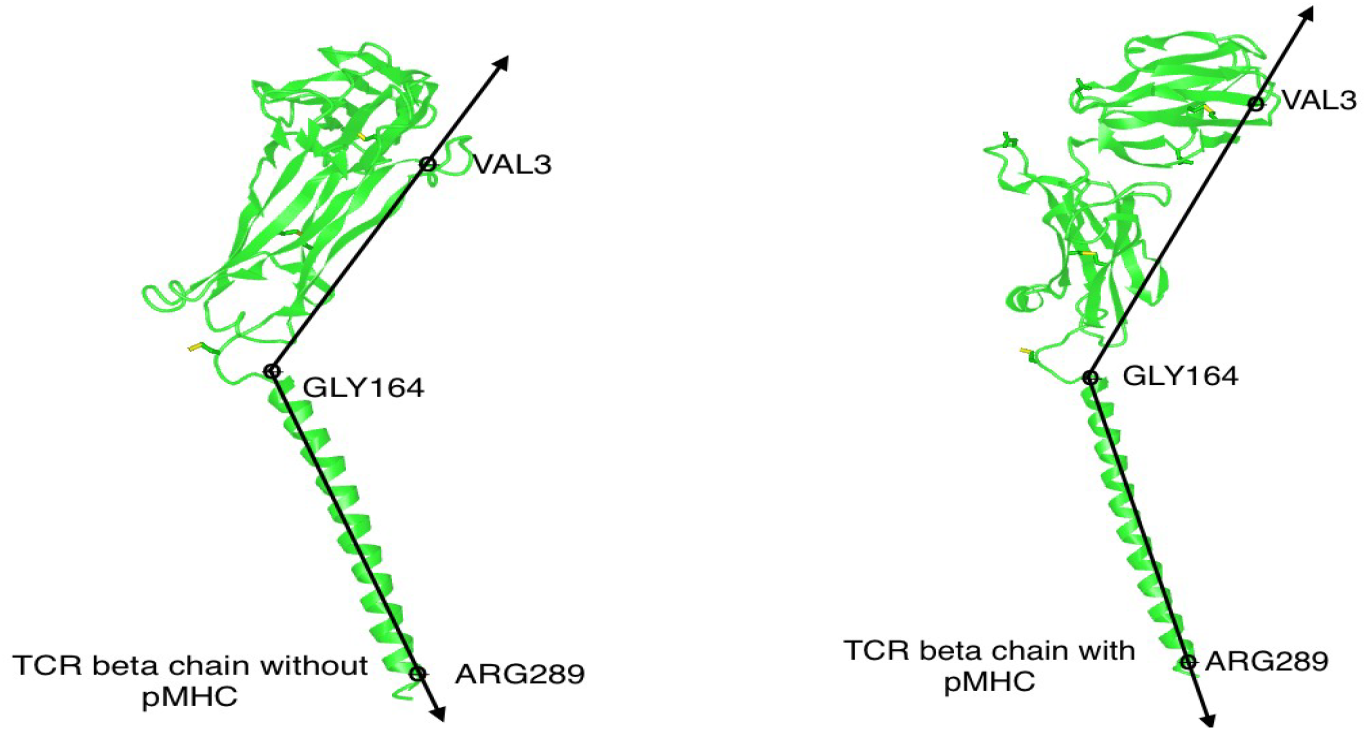
Visualization of the atoms used to calculate the tilt angle, specifically TCR*β* residues VAL_3_, GLY_164_, and ARG_289_. The left panel shows the TCR*β* chain in its unbound state, while the right panel displays the TCR*β* chain in complex with pMHC, highlighting a shift in the tilt angle from 104.99^°^ to 147.56^°^ upon ligand binding.

**Fig. B2.**
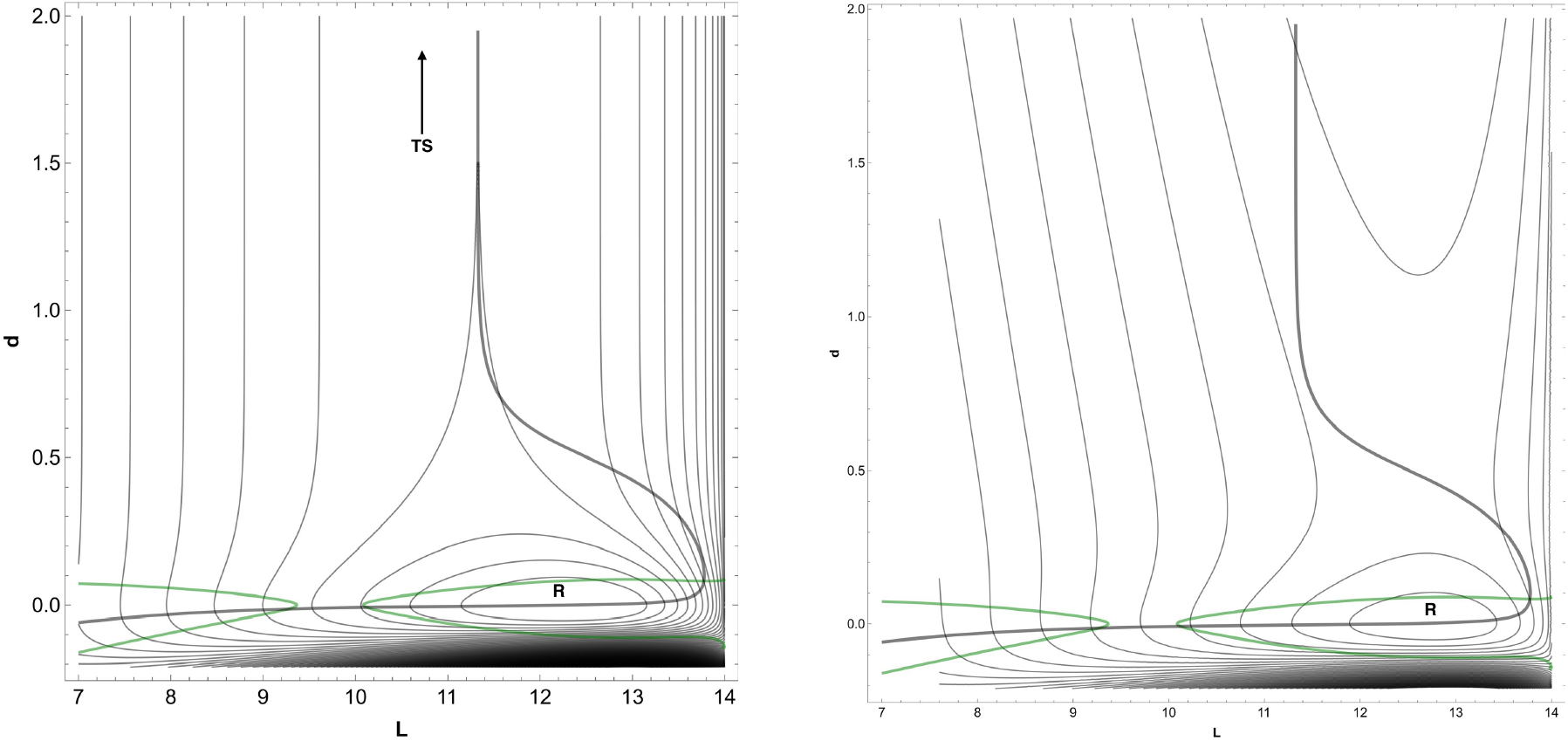
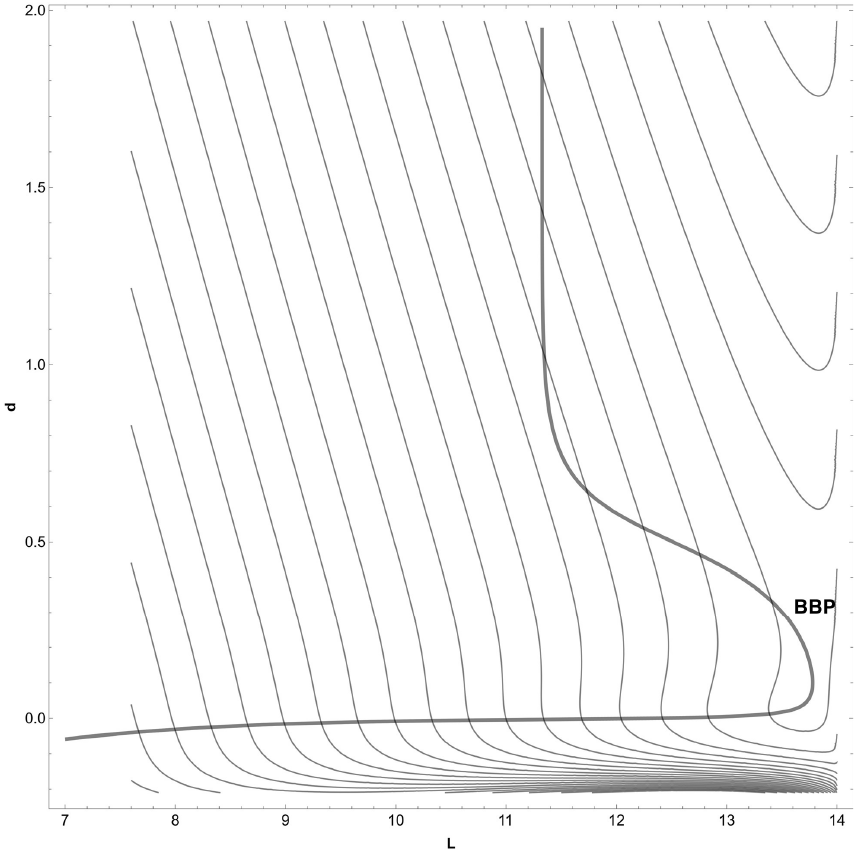
2D potential energy surface (PES) for TCR–antagonist–pMHC interaction under high force (*F* = 75), showing deformation of the energy landscape with applied load.

## References

[1] Markus G Rudolph, Robyn L Stanfield, and Ian A Wilson. How TCRs bind MHCs, peptides, and coreceptors. Annu. Rev. Immunol., 24(1):419–466, 2006.

[2] Eric J Sundberg and Roy A Mariuzza. Molecular recognition in antibody-antigen complexes. Advances in protein chemistry, 61:119–160, 2002.

[3] Jamie Rossjohn, Stephanie Gras, John J Miles, Stephen J Turner, Dale I Godfrey, and James McCluskey. T cell antigen receptor recognition of antigen-presenting molecules. Annual review of immunology, 33(1):169–200, 2015.

[4] Jun Huang, Mario Brameshuber, Xun Zeng, Jianming Xie, Qi-jing Li, Yueh-hsiu Chien, Salvatore Valitutti, and Mark M Davis. A single peptide-major histocompatibility complex ligand triggers digital cytokine secretion in CD4+ T cells. Immunity, 39(5):846–857, 2013.

[5] Kristine N Brazin, Robert J Mallis, Dibyendu Kumar Das, Yinnian Feng, Wonmuk Hwang, Jia-huai Wang, Gerhard Wagner, Matthew J Lang, and Ellis L Reinherz. Structural features of the ??βTCR mechanotransduction apparatus that promote pMHC discrimination. Frontiers in immunology, 6:441, 2015.

[6] Michel A Cuendet and Olivier Michielin. Protein-protein interaction investigated by steered molecular dynamics: the TCR-pMHC complex. Biophysical journal, 95(8):3575–3590, 2008.

[7] Kathryn M Armstrong, Kurt H Piepenbrink, and Brian M Baker. Conformational changes and flexibility in T-cell receptor recognition of peptide–MHC complexes. Biochemical Journal, 415(2):183–196, 2008.

[8] Sun Taek Kim, Koh Takeuchi, Zhen-Yu J Sun, Maki Touma, Carlos E Castro, Amr Fahmy, Matthew J Lang, Gerhard Wagner, and Ellis L Reinherz. The ??β T cell receptor is an anisotropic mechanosensor. Journal of Biological Chemistry, 284(45):31028–31037, 2009.

[9] Mark S Lee, Caleb R Glassman, Neha R Deshpande, Hemant B Badgandi, Heather L Parrish, Chayasith Uttamapinant, Philipp S Stawski, Alice Y Ting, and Michael S Kuhns. A mechanical switch couples T cell receptor triggering to the cytoplasmic juxtamembrane regions of CD3ζ ζ. Immunity, 43(2):227–239, 2015.

[10] Baoyu Liu, Wei Chen, Brian D Evavold, and Cheng Zhu. Accumulation of dynamic catch bonds between TCR and agonist peptide-MHC triggers T cell signaling. Cell, 157(2): 357–368, 2014.

[11] Rong Ma, Anna V Kellner, Victor Pui-Yan Ma, Hanquan Su, Brendan R Deal, Joshua M Brockman, and Khalid Salaita. DNA probes that store mechanical information reveal transient piconewton forces applied by T cells. Proceedings of the National Academy of Sciences, 116(34):16949–16954, 2019.

[12] Yang Liu, Lori Blanchfield, Victor Pui-Yan Ma, Rakieb Andargachew, Kornelia Galior, Zheng Liu, Brian Evavold, and Khalid Salaita. DNA-based nanoparticle tension sensors reveal that T-cell receptors transmit defined pN forces to their antigens for enhanced fidelity. Proceedings of the National Academy of Sciences, 113(20):5610–5615, 2016.

[13] Hyun-Kyu Choi, Peiwen Cong, Chenghao Ge, Aswin Natarajan, Baoyu Liu, Yong Zhang, Kaitao Li, Muaz Nik Rushdi, Wei Chen, Jizhong Lou, et al. Catch bond models may explain how force amplifies TCR signaling and antigen discrimination. Nature communications, 14(1):2616, 2023.

[14] Yinnian Feng, Kristine N Brazin, Eiji Kobayashi, Robert J Mallis, Ellis L Reinherz, and Matthew J Lang. Mechanosensing drives acuity of ??β T-cell recognition. Proceedings of the National Academy of Sciences, 114(39):E8204–E8213, 2017.

[15] Kristin A Hogquist, Stephen C Jameson, William R Heath, Jane L Howard, Michael J Bevan, and Francis R Carbone. T cell receptor antagonist peptides induce positive selection. Cell, 76(1):17–27, 1994.

[16] Leah V Sibener, Ricardo A Fernandes, Elizabeth M Kolawole, Catherine B Carbone, Fan Liu, Darren McAffee, Michael E Birnbaum, Xinbo Yang, Laura F Su, Wong Yu, et al. Isolation of a structural mechanism for uncoupling T cell receptor signaling from peptide-MHC binding. Cell, 174(3):672–687, 2018.

[17] Pierre-Henri Puech and Pierre Bongrand. Mechanotransduction as a major driver of cell behaviour: Mechanisms, and relevance to cell organization and future research. Open Biology, 11(11):210256, 2021.

[18] Evgeni V Sokurenko, Viola Vogel, and Wendy E Thomas. Catch-bond mechanism of force-enhanced adhesion: counterintuitive, elusive, but… widespread? Cell host & microbe, 4 (4):314–323, 2008.

[19] Kevin Y Chen, Edward Jenkins, Markus Körbel, Aleks Ponjavic, Anna H Lippert, Ana Mafalda Santos, Nicole Ashman, Caitlin O’Brien-Ball, Jemma McBride, David Klenerman, et al. Trapping or slowing the diffusion of T cell receptors at close contacts initiates T cell signaling. Proceedings of the National Academy of Sciences, 118(39):e2024250118, 2021.

[20] Anna-Lisa Lanz, Giulia Masi, Nicla Porciello, André Cohnen Deborah Cipria, Dheeraj Prakaash, Štefan Bálint, Roberto Raggiaschi, Donatella Galgano, David K Cole, et al. Allosteric activation of T cell antigen receptor signaling by quaternary structure relaxation. Cell reports, 36(2), 2021.

[21] Kannan Natarajan, Andrew C McShan, Jiansheng Jiang, Vlad K Kumirov, Rui Wang, Huaying Zhao, Peter Schuck, Mulualem E Tilahun, Lisa F Boyd, Jinfa Ying, et al. An allosteric site in the T-cell receptor Cβ domain plays a critical signalling role. Nature communications, 8(1):15260, 2017.

[22] Yanan He, Pragati Agnihotri, Sneha Rangarajan, Yihong Chen, Melissa C Kerzic, Buyong Ma, Ruth Nussinov, Roy A Mariuzza, and John Orban. Peptide–MHC binding reveals conserved allosteric sites in MHC class I-and class II-restricted T cell receptors (TCRs). Journal of molecular biology, 432(24):166697, 2020.

[23] Jinsung Hong, Chenghao Ge, Prithiviraj Jothikumar, Zhou Yuan, Baoyu Liu, Ke Bai, Kaitao Li, William Rittase, Miho Shinzawa, Yun Zhang, et al. A TCR mechanotransduction signaling loop induces negative selection in the thymus. Nature immunology, 19(12):1379– 1390, 2018.

[24] Ondrej Stepanek, Arvind S Prabhakar, Celine Osswald, Carolyn G King, Anna Bulek, Dieter Naeher, Marina Beaufils-Hugot, Michael L Abanto, Virginie Galati, Barbara Hausmann, et al. Coreceptor scanning by the T cell receptor provides a mechanism for T cell tolerance. Cell, 159(2):333–345, 2014.

[25] Timothy K Starr, Stephen C Jameson, and Kristin A Hogquist. Positive and negative selection of T cells. Annual review of immunology, 21(1):139–176, 2003.

[26] Laura M DeFord-Watts, Tara C Tassin, Amy M Becker, Jennifer J Medeiros, Joseph P Albanesi, Paul E Love, Christoph Wulfing, and Nicolai SC Van Oers. The cytoplasmic tail of the T cell receptor CD3 ε subunit contains a phospholipid-binding motif that regulates T cell functions. The Journal of Immunology, 183(2):1055–1064, 2009.

[27] Chenqi Xu, Etienne Gagnon, Matthew E Call, Jason R Schnell, Charles D Schwieters, Christopher V Carman, James J Chou, and Kai W Wucherpfennig. Regulation of T cell receptor activation by dynamic membrane binding of the CD3 ε cytoplasmic tyrosine-based motif. Cell, 135(4):702–713, 2008.

[28] Hao Zhang, Shaun-Paul Cordoba, Omer Dushek, and P Anton Van Der Merwe. Basic residues in the T-cell receptor ζ cytoplasmic domain mediate membrane association and modulate signaling. Proceedings of the National Academy of Sciences, 108(48):19323– 19328, 2011.

[29] Luis M Fernández-Aguilar, Inmaculada Vico-Barranco, Mikel M Arbulo-Echevarria, and Enrique Aguado. A story of kinases and adaptors: The role of LCK, ZAP-70 and LAT in switch panel governing T-cell development and activation. Biology, 12(9):1163, 2023.

[30] Jeanne C Stachowiak, Eva M Schmid, Christopher J Ryan, Hyoung Sook Ann, Darryl Y Sasaki, Michael B Sherman, Phillip L Geissler, Daniel A Fletcher, and Carl C Hayden. Membrane bending by protein–protein crowding. Nature cell biology, 14(9):944–949, 2012.

[31] Jeanne C Stachowiak, Frances M Brodsky, and Elizabeth A Miller. A cost–benefit analysis of the physical mechanisms of membrane curvature. Nature cell biology, 15(9):1019–1027, 2013.

[32] Ozlem Keskin. Binding induced conformational changes of proteins correlate with their intrinsic fluctuations: a case study of antibodies. BMC structural biology, 7:1–11, 2007.

[33] P Anton van der Merwe and Simon J Davis. Molecular interactions mediating T cell antigen recognition. Annual review of immunology, 21(1):659–684, 2003.

[34] Roy A Mariuzza, Pragati Agnihotri, and John Orban. The structural basis of T-cell receptor (TCR) activation: An enduring enigma. Journal of Biological Chemistry, 295(4):914–925, 2020.

[35] Ian A Wilson and K Christopher Garcia. T-cell receptor structure and TCR complexes. Current Opinion in Structural Biology, 7(6):839–848, 1997.

[36] Antonio de la Hera, U Müller, Carina Olsson, Sylvie Isaaz, and Alan Tunnacliffe. Structure of the T cell antigen receptor (TCR): two CD3 epsilon subunits in a functional TCR/CD3 complex. The Journal of experimental medicine, 173(1):7–17, 1991.

[37] Mohammad Ameen Al-Aghbar, Ashwin K Jainarayanan, Michael L Dustin, and Steve R Roffler. The interplay between membrane topology and mechanical forces in regulating T cell receptor activity. Communications Biology, 5(1):40, 2022.

[38] Floris J van Eerden, Aalaa Alrahman Sherif, Mara Anais Llamas-Covarrubias, Arthur Millius, Xiuyuan Lu, Shigenari Ishizuka, Sho Yamasaki, and Daron M Standley. TCRpMHC complex formation triggers CD3 dynamics. eLife, 12, 2023.

[39] Prithvi R Pandey, Bartosz Różycki, Reinhard Lipowsky, and Thomas R Weikl. Structural variability and concerted motions of the T cell receptor–CD3 complex. Elife, 10:e67195, 2021.

[40] De Dong, Lvqin Zheng, Jianquan Lin, Bailing Zhang, Yuwei Zhu, Ningning Li, Shuangyu Xie, Yuhang Wang, Ning Gao, and Zhiwei Huang. Structural basis of assembly of the human T cell receptor–CD3 complex. Nature, 573(7775):546–552, 2019.

[41] Ellis L Reinherz. The structure of a T-cell mechanosensor, 2019.

[42] Yanan He, Sneha Rangarajan, Melissa Kerzic, Ming Luo, Yihong Chen, Qian Wang, Yiyuan Yin, Creg J Workman, Kate M Vignali, Dario AA Vignali, et al. Identification of the docking site for CD3 on the T cell receptor β chain by solution NMR. Journal of Biological Chemistry, 290(32):19796–19805, 2015.

[43] Dheeraj Prakaash, Graham P Cook, Oreste Acuto, and Antreas C Kalli. Multi-scale simulations of the T cell receptor reveal its lipid interactions, dynamics and the arrangement of its cytoplasmic region. PLOS Computational Biology, 17(7):e1009232, 2021.

[44] Casey O Barkan and Robijn F Bruinsma. Topology of molecular deformations induces triphasic catch bonding in selectin–ligand bonds. Proceedings of the National Academy of Sciences, 121(6):e2315866121, 2024.

[45] Kei Saotome, Drew Dudgeon, Kiersten Colotti, Michael J Moore, Jennifer Jones, Yi Zhou, Ashique Rafique, George D Yancopoulos, Andrew J Murphy, John C Lin, et al. Structural analysis of cancer-relevant TCR-CD3 and peptide-MHC complexes by cryoEM. Nature Communications, 14(1):2401, 2023.

[46] Wolfgang Quapp, Josep Maria Bofill, and Jordi Ribas-Ariño. Toward a theory of mechanochemistry: Simple models from the very beginnings. International Journal of Quantum Chemistry, 118(23):e25775, 2018.

[47] Wolfgang Quapp and Josep Maria Bofill. Mechanochemistry on the m üller–b rown surface by Newton trajectories. International Journal of Quantum Chemistry, 118(8): e25522, 2018.

[48] Wolfgang Quapp and Josep Maria Bofill. Theory and Examples of Catch Bonds. The Journal of Physical Chemistry B, 128(17):4097–4110, 2024.

[49] Wolfgang Quapp and Josep Maria Bofill. Reaction rates in a theory of mechanochemical pathways. Journal of computational chemistry, 37(27):2467–2478, 2016.

[50] Yohichi Suzuki and Olga K Dudko. Biomolecules under mechanical stress: a simple mechanism of complex behavior. The Journal of chemical physics, 134(6), 2011.

[51] Sai Sriharsha M Konda, Johnathan N Brantley, Bibin T Varghese, Kelly M Wiggins, Christopher W Bielawski, and Dmitrii E Makarov. Molecular catch bonds and the anti-hammond effect in polymer mechanochemistry. Journal of the American Chemical Society, 135(34):12722–12729, 2013.

[52] Wendy E Thomas. Understanding the counterintuitive phenomenon of catch bonds. Current Nanoscience, 3(1):63–77, 2007.

[53] Dibyendu Kumar Das, Yinnian Feng, Robert J Mallis, Xiaolong Li, Derin B Keskin, Rebecca E Hussey, Sonia K Brady, Jia-Huai Wang, Gerhard Wagner, Ellis L Reinherz, et al. Force-dependent transition in the T-cell receptor β-subunit allosterically regulates peptide discrimination and pMHC bond lifetime. Proceedings of the National Academy of Sciences, 112(5):1517–1522, 2015.

[54] Peng Wu, Tongtong Zhang, Baoyu Liu, Panyu Fei, Lei Cui, Rui Qin, Huaying Zhu, Danmei Yao, Ryan J Martinez, Wei Hu, et al. Mechano-regulation of peptide-MHC class I conformations determines TCR antigen recognition. Molecular cell, 73(5):1015–1027, 2019.

[55] Lukas Sušac, Mai T Vuong, Christoph Thomas, Sören von Bülow, Caitlin O’Brien-Ball, Ana Mafalda Santos, Ricardo A Fernandes, Gerhard Hummer, Robert Tampé, and Simon J Davis. Structure of a fully assembled tumor-specific T cell receptor ligated by pMHC. Cell, 185(17):3201–3213, 2022.

[56] Wolfgang Quapp, Josep Maria Bofill, and Jordi Ribas-Ariño. Analysis of the Acting Forces in a Theory of Catalysis and Mechanochemistry. The Journal of Physical Chemistry A, 121(14):2820–2838, 2017.

[57] Hrant P Hratchian and H Bernhard Schlegel. Finding minima, transition states, and following reaction pathways on ab initio potential energy surfaces. In Theory and applications of computational chemistry, pages 195–249. Elsevier, 2005.

[58] Shishir Adhikari, Jacob Moran, Christopher Weddle, and Michael Hinczewski. Unraveling the mechanism of the cadherin-catenin-actin catch bond. PLoS computational biology, 14 (8):e1006399, 2018.

[59] Shaon Chakrabarti, Michael Hinczewski, and D Thirumalai. Plasticity of hydrogen bond networks regulates mechanochemistry of cell adhesion complexes. Proceedings of the National Academy of Sciences, 111(25):9048–9053, 2014.

[60] Evan Evans and Ken Ritchie. Dynamic strength of molecular adhesion bonds. Biophysical journal, 72(4):1541–1555, 1997.

[61] Wendy E Thomas, Viola Vogel, and Evgeni Sokurenko. Biophysics of catch bonds. Annu. Rev. Biophys., 37(1):399–416, 2008.

[62] Franklin H Branin. Widely convergent method for finding multiple solutions of simultaneous nonlinear equations. IBM Journal of Research and Development, 16(5):504–522, 1972.

[63] Immo Diener and Robert Schaback. An extended continuous Newton method. Journal of Optimization Theory and Applications, 67:57–77, 1990.

[64] Wolfgang Quapp, Michael Hirsch, Olaf Imig, and Dietmar Heidrich. Searching for saddle points of potential energy surfaces by following a reduced gradient. Journal of computational chemistry, 19(9):1087–1100, 1998.

[65] Wolfgang Quapp, Michael Hirsch, and Dietmar Heidrich. Bifurcation of reaction pathways: the set of valley ridge inflection points of a simple three-dimensional potential energy surface. Theoretical Chemistry Accounts, 100:285–299, 1998.

[66] Michael Hirsch and Wolfgang Quapp. The reaction pathway of a potential energy surface as curve with induced tangent. Chemical physics letters, 395(1-3):150–156, 2004.

[67] George I Bell. Models for the specific adhesion of cells to cells: a theoretical framework for adhesion mediated by reversible bonds between cell surface molecules. Science, 200 (4342):618–627, 1978.

[68] Jonathon Howard and RL Clark. Mechanics of motor proteins and the cytoskeleton. Appl. Mech. Rev., 55(2):B39–B39, 2002.

[69] Philip M Morse. Diatomic molecules according to the wave mechanics II. vibrational levels. Physical review, 34(1):57, 1929.

[70] Terrell L Hill. Free energy transduction and biochemical cycle kinetics. Courier Corporation, 2005.

[71] Roger A Horn and Charles R Johnson. Matrix analysis. Cambridge university press, 2012.

[72] Shaomin Tian, Robert Maile, Edward J Collins, and Jeffrey A Frelinger. CD8+ T cell activation is governed by TCR-peptide/MHC affinity, not dissociation rate. The Journal of Immunology, 179(5):2952–2960, 2007.

[73] S Munir Alam, G Mark Davies, Christina M Lin, Tomasz Zal, Wade Nasholds, Stephen C Jameson, Kristin A Hogquist, Nicholas RJ Gascoigne, and Paul J Travers. Qualitative and quantitative differences in T cell receptor binding of agonist and antagonist ligands. Immunity, 10(2):227–237, 1999.

[74] Jonathan M Boulter, Nicole Schmitz, Andrew K Sewell, Andrew J Godkin, Martin F Bachmann, and Awen M Gallimore. Potent T cell agonism mediated by a very rapid TCR/pMHC interaction. European journal of immunology, 37(3):798–806, 2007.

[75] Terrell L Hill. Cooperativity theory in biochemistry: steady-state and equilibrium systems. Springer Science & Business Media, 2013.

[76] Ken A Dill. Dominant forces in protein folding. Biochemistry, 29(31):7133–7155, 1990.

[77] Michael K Gilson, James A Given, Bruce L Bush, and J Andrew McCammon. The statistical-thermodynamic basis for computation of binding affinities: a critical review. Biophysical journal, 72(3):1047–1069, 1997.

[78] Jun Huang, Veronika I Zarnitsyna, Baoyu Liu, Lindsay J Edwards, Ning Jiang, Brian D Evavold, and Cheng Zhu. The kinetics of two-dimensional TCR and pMHC interactions determine T-cell responsiveness. Nature, 464(7290):932–936, 2010.

[79] Eric F Pettersen, Thomas D Goddard, Conrad C Huang, Gregory S Couch, Daniel M Greenblatt, Elaine C Meng, and Thomas E Ferrin. UCSF Chimera—a visualization system for exploratory research and analysis. Journal of computational chemistry, 25(13): 1605–1612, 2004.

[80] National Library of Medicine. Molecular modeling database (mmdb), 2024. Accessed via https://www.ncbi.nlm.nih.gov/Structure/.

[81] DV Shevyrev, VP Tereshchenko, and SV Sennikov. The enigmatic nature of the TCRpMHC interaction: implications for CAR-T and TCR-T engineering. International Journal of Molecular Sciences, 23(23):14728, 2022.

[82] J. N. Langley. On the reaction of cells and of nerve endings to certain poisons. Journal of Physiology, 33(4-5):374–413, 1905.

[83] A. J. Clark. The reaction between acetylcholine and muscle cells. Journal of Physiology, 61(4):530–546, 1933.

[84] R. P. Stephenson. A modification of receptor theory. British Journal of Pharmacology and Chemotherapy, 11(4):379–393, 1956.

[85] E.J. Ariëns. Affinity and intrinsic activity in the theory of competitive inhibition. Archives Internationales de Pharmacodynamie et de Thérapie, 99:32–49, 1954.

[86] J. H. Gaddum. The quantitative effects of antagonistic drugs. Journal of Physiology, 89: 7P–9P, 1937.

[87] Svante Arrhenius. Über die Reaktionsgeschwindigkeit bei der Inversion von Rohrzucker durch Säuren. Zeitschrift für physikalische Chemie, 4(1):226–248, 1889.

[88] Hendrik Anthony Kramers. Brownian motion in a field of force and the diffusion model of chemical reactions. physica, 7(4):284–304, 1940.

[89] Peter Hänggi, Peter Talkner, and Michal Borkovec. Reaction-rate theory: fifty years after Kramers. Reviews of modern physics, 62(2):251, 1990.

[90] James S Langer. Statistical theory of the decay of metastable states. Annals of Physics, 54 (2):258–275, 1969.

