## Supplementary material for "Mechanochemical Energy Landscapes Under Force: Catch–Slip Bonds in T-Cell Activation"

### Model Equations

We study the total free energy $V_f$ of a TCR–pMHC complex as a function of

$$L \in [0, 2W], \qquad d \geq 0,$$

where $L$ is the protein extension and $d$ is the ligand-binding site separation. The hinge angle $\theta$ is related to $L$ by

$$\theta(L) = 2 \arcsin\left(\frac{L}{2W}\right), \tag{1}$$

$$L(\theta) = 2W \sin\left(\frac{\theta}{2}\right). \tag{2}$$

The half-angle identity

$$\sqrt{1 - \frac{L^2}{4W^2}} \;=\; \cos\big(\theta(L)/2\big) \qquad \Rightarrow \qquad \frac{\mathrm{d}\theta}{\mathrm{d}L} \;=\; \frac{1}{W\sqrt{1 - L^2/(4W^2)}} \;=\; \frac{1}{W \cos(\theta/2)} \tag{3}$$

will be used below.

The total free energy is

$$V_f(L, d) = V(\theta(L)) + D(\theta(L))\,\Phi(d) - F\,(L + d), \tag{4}$$

where $F \geq 0$ is an external force, the hinge energy is

$$V(\theta) = \frac{1}{2} k_\theta (\theta - \theta_0)^2, \qquad k_\theta > 0, \tag{5}$$

and the (Morse) factor is written via $u := e^{-d/d_0} \in (0, 1]$:

$$\Phi(d) = \left(1 - e^{-d/d_0}\right)^2 - 1 = u^2 - 2u.$$

The depth modulation is Gaussian:

$$D(\theta) = D_0 \exp\left[-\frac{(\theta-\theta_1)^2}{2\sigma^2}\right], \qquad D_0 > 0,\ \sigma > 0. \tag{6}$$

### 1 Existence and multiplicity of steady states

#### Stationarity in $d$: Equation (A)

Differentiating $\Phi$ via $u = e^{-d/d_0}$ with $\mathrm{d}u/\mathrm{d}d = -u/d_0$ gives

$$\frac{\mathrm{d}\Phi}{\mathrm{d}d} = \frac{\mathrm{d}}{\mathrm{d}d}(u^2 - 2u) = 2u\frac{\mathrm{d}u}{\mathrm{d}d} - 2\frac{\mathrm{d}u}{\mathrm{d}d} = \frac{2}{d_0}u(1-u).$$

Hence the $d$-stationarity condition is

$$\boxed{\frac{2}{d_0}D(\theta)\,u(1-u) \;=\; F.} \tag{A}$$

Introduce

$$\alpha := \frac{Fd_0}{2} > 0, \qquad C(\theta) := \frac{\alpha}{D(\theta)} > 0, \tag{7}$$

so (A) is equivalent to

$$\boxed{u^2 - u + C(\theta) = 0, \qquad u \in (0,1].} \tag{8}$$

The discriminant $\Delta(\theta) = 1 - 4C(\theta)$ yields the admissibility threshold

$$\boxed{D(\theta) \geq 2Fd_0} \tag{9}$$

with roots

$$\boxed{u_\pm(\theta) = \frac{1 \pm \sqrt{1-4C(\theta)}}{2}.} \tag{10}$$

Using $D(\theta) = D_0\exp(-(\theta-\theta_1)^2/(2\sigma^2))$, (9) gives the admissible angle band

$$\boxed{\theta \in [\theta_-, \theta_+], \quad \theta_\pm := \theta_1 \pm \delta, \quad \delta := \sigma\sqrt{2\log\left(\frac{D_0}{2Fd_0}\right)},} \tag{11}$$

nonempty iff $D_0 \geq 2Fd_0$.

From (8), $u^2 = u - C(\theta)$, hence

$$\boxed{\Phi(d) = u^2 - 2u = -C(\theta) - u \;=\; -\frac{\alpha}{D(\theta)} - u.} \tag{$\star$}$$

#### Stationarity in $L$: Equation (B)

Differentiate $V_f$ with respect to $L$, noting that $d$ is independent of $L$:

$$\frac{\partial V_f}{\partial L} = \left[\,V'(\theta) + D'(\theta)\,\Phi(d)\,\right]\frac{\mathrm{d}\theta}{\mathrm{d}L} \;-\; F.$$

Using $V'(\theta) = k_\theta(\theta - \theta_0)$, $D'(\theta) = -\frac{\theta-\theta_1}{\sigma^2}D(\theta)$, and (22), we obtain

$$\left[k_\theta(\theta - \theta_0) - \frac{\theta - \theta_1}{\sigma^2}D(\theta)\,\Phi(d)\right] \cdot \frac{1}{W\cos(\theta/2)} = F. \tag{B}$$

Multiplying by $W\cos(\theta/2)$ gives the affine form

$$k_\theta(\theta - \theta_0) - \frac{\theta - \theta_1}{\sigma^2}D(\theta)\,\Phi(d) = FW\cos(\theta/2) =: R(\theta). \tag{B1}$$

#### Linearizing (B) in $u$ and reduction to one scalar

Define

$$A(\theta) := k_\theta(\theta - \theta_0) + \frac{\alpha}{\sigma^2}(\theta - \theta_1), \qquad B(\theta) := \frac{\theta - \theta_1}{\sigma^2}D(\theta), \qquad R(\theta) := FW\cos(\theta/2).$$

Substituting $(\star)$ into $(\mathrm{B1})$ yields the linear relation

$$\boxed{A(\theta) + B(\theta)\,u = R(\theta).} \tag{B-linear}$$

If $B(\theta) \neq 0$, then $u = (R - A)/B$. Enforcing (A) gives

$$\left(\frac{R-A}{B}\right)\left(1 - \frac{R-A}{B}\right) = C(\theta) \iff (R-A)B - (R-A)^2 - C(\theta)B^2 = 0.$$

Hence the system reduces to

$$\boxed{J(\theta) := (R-A)\,B - (R-A)^2 - C(\theta)\,B^2 = 0} \quad \text{on} \quad \Theta = \{\theta : D(\theta) \geq 2Fd_0\}. \tag{12}$$

#### How $J(\theta) = 0$ interacts with the two roots of (A)

For $\theta \in \Theta$, (A) has roots $u_\pm(\theta)$ in (10) (double root $u = \frac{1}{2}$ at $\theta_\pm$). For $\theta \neq \theta_1$, (B-linear) gives $u = (R-A)/B$, so $J(\theta) = 0$ iff that $u$ satisfies $u(1-u) = C(\theta)$. Each root $\theta^* \in \Theta$ of $J$ therefore selects a unique $u^* \in (0, 1]$ solving both equations, with

$$d^* = -d_0 \ln u^*, \qquad L^* = 2W\sin\left(\frac{\theta^*}{2}\right).$$

At the center $\theta = \theta_1$, $B = 0$ and (B-linear) reduces to

$$A(\theta_1) = R(\theta_1) \iff k_\theta(\theta_1 - \theta_0) = FW\cos(\theta_1/2),$$

and $J(\theta_1) = -(R-A)^2 \leq 0$ with equality exactly under this condition. If additionally $D(\theta_1) > 2Fd_0$, both $u_\pm(\theta_1) \in (0, 1)$ are admissible, giving two separations at the same angle $\theta_1$. At $\theta_\pm$, $D = 2Fd_0$ and $u = \frac{1}{2}$; substituting gives

$$J(\theta_\pm) = -\Big(R(\theta_\pm) - A(\theta_\pm) \mp \tfrac{1}{2}\,B(\theta_\pm)\Big)^2 \leq 0.$$

#### Unimodality of $J(\theta)$ on each side of $\theta_1$

Recall

$$J(\theta) = -G(\theta)^2 + G(\theta)\,B(\theta) - C(\theta)B(\theta)^2, \quad G := R - A,$$

and the concave-envelope bound

$$J(\theta) \;\le\; J_{\max}(\theta) := B(\theta)^2\Big(\tfrac{1}{4} - C(\theta)\Big), \tag{13}$$

with $J_{\max}(\theta) > 0$ iff $D(\theta) > 2Fd_0$. Writing $\varepsilon := \theta - \theta_1$,

$$J_{\max}(\theta) = \underbrace{\Big(\tfrac{\varepsilon^2}{\sigma^4} e^{-\varepsilon^2/(2\sigma^2)}\Big)}_{=:h_1(\varepsilon)} \cdot \underbrace{D_0\Big(\tfrac{D(\theta)}{4} - \alpha\Big)}_{=:h_2(\varepsilon)}, \tag{14}$$

where $h_1$ is single-hump and $h_2$ is strictly decreasing in $|\varepsilon|$, vanishing at $|\varepsilon| = \delta$ (where $D = 2Fd_0$). The product is therefore unimodal per side; consequently $J$ can cross the axis at most once on each side of $\theta_1$.

**Proposition 1.1** (Exactly one steady state: degenerate cases only)**.** *Let $J(\theta) = (R-A)B-(R-A)^2 - C(\theta)B^2$ on the admissible band $\Theta = \{\theta : D(\theta) \ge 2Fd_0\}$ (equivalently, $\Theta = [\theta_-, \theta_+]$ when $D_0 \ge 2Fd_0$). The full system has* exactly one *steady state if and only if one of the following holds:*

*(C)* ***Center–threshold degeneracy:***

$$A(\theta_1) = R(\theta_1) \quad \textit{and} \quad D(\theta_1) = 2Fd_0,$$

*i.e. $k_\theta(\theta_1 - \theta_0) = FW\cos(\theta_1/2)$ and Equation (A) has the double root $u = \frac{1}{2}$. Outcome: a unique steady state at $\theta = \theta_1$.*

*(S)* ***Single-side tangency:*** *There exists $\theta^* \in (\theta_-, \theta_1) \cup (\theta_1, \theta_+)$ such that*

$$J(\theta^*) = 0, \qquad J'(\theta^*) = 0, \qquad J''(\theta^*) < 0,$$

*i.e. $J$ touches the axis once on one side and has no other zeros. Outcome: a unique off-center steady state.*

*In all other admissible (nondegenerate) regimes, the system has* 0 *or* 2 *steady states (never more than two). Both* (C) *and* (S) *are codimension-one.*

#### 2 Stability analysis

From our total free energy $V_f$ of a TCR-pMHC complex:

$$L \in [0, 2W], \qquad d \geq 0,$$

where $L$ is the protein extension and $d$ is the ligand-binding site separation. The hinge angle $\theta$ is related to $L$ by

$$\theta(L) = 2 \arcsin\left(\frac{L}{2W}\right), \tag{15}$$

$$L(\theta) = 2W \sin\left(\frac{\theta}{2}\right), \tag{16}$$

with $W > 0$.

**Total free energy.**

$$V_f(L, d) \;=\; V\big(\theta(L)\big) \;+\; B\big(d, \theta(L)\big) \;-\; F\,(L + d), \tag{17}$$

where $F \geq 0$ is an external force.

**Hinge (angular) energy.**

$$V(\theta(L)) \;=\; \tfrac{1}{2}\, k_\theta \left(\theta(L) - \theta_0\right)^2, \qquad k_\theta > 0. \tag{18}$$

**Ligand binding energy (Morse form with angle-dependent depth).**

$$B(d, \theta(L)) = D\big(\theta(L)\big)\, \Phi(d), \tag{19}$$

$$\Phi(d) := \left(1 - e^{-d/d_0}\right)^2 - 1, \qquad d_0 > 0, \tag{20}$$

with angle-dependent depth

$$D\big(\theta(L)\big) \;=\; D_0 \exp\left[-\frac{(\theta(L) - \theta_1)^2}{2\sigma^2}\right], \qquad D_0 > 0,\; \sigma > 0. \tag{21}$$

For convenience, define

$$u := e^{-d/d_0} \in (0, 1], \qquad \Phi(d) = u^2 - 2u.$$

**Elementary derivatives we will need.**

**(Angle geometry)** $$\frac{d\theta(L)}{dL} = 2 \cdot \frac{1}{\sqrt{1-(L/2W)^2}} \cdot \frac{1}{2W} = \frac{1}{W\sqrt{1-\frac{L^2}{4W^2}}}. \tag{22}$$

**(Morse factor)** $$\frac{du}{dd} = -\frac{u}{d_0}, \qquad \frac{d\Phi}{dd} = \frac{d}{dd}\big(u^2 - 2u\big) = 2u\,\frac{du}{dd} - 2\,\frac{du}{dd} = \frac{2}{d_0}\,u(1-u). \tag{23}$$

**(Depth $D$)** $$\underbrace{\frac{d}{d\theta}D(\theta)}_{=:D'(\theta)} = -\frac{\theta-\theta_1}{\sigma^2}\,D(\theta), \qquad \underbrace{\frac{d}{dL}D\big(\theta(L)\big)}_{=\,D'(\theta(L))\,\theta'(L)} = D\big(\theta(L)\big)\,\theta'(L)\left(-\frac{\theta(L)-\theta_1}{\sigma^2}\right). \tag{24}$$

### Derivative with respect to $d$

Starting from (17), using (19) and (23):

$$\frac{\partial V_f}{\partial d} = \frac{\partial}{\partial d}\Big[D\big(\theta(L)\big)\,\Phi(d)\Big] - F.$$

Because $D(\theta(L))$ does *not* depend on $d$, we have

$$\frac{\partial V_f}{\partial d} = D\big(\theta(L)\big)\,\frac{d\Phi}{dd} - F.$$

Insert (23):

$$\begin{aligned}\frac{\partial V_f}{\partial d} &= D\big(\theta(L)\big)\left(\frac{2}{d_0}\,u(1-u)\right) - F \\ &= \boxed{\frac{2\,D\big(\theta(L)\big)}{d_0}\,u(1-u) - F}.\end{aligned} \tag{25}$$

*Critical-point equation in $d$.* Set (25) to zero and write $D := D(\theta(L))$ for brevity:

$$\frac{2D}{d_0}\,u(1-u) - F = 0 \iff u(1-u) = \frac{Fd_0}{2D}. \tag{26}$$

Expanding the left-hand side $u(1-u) = u - u^2$ yields

$$u^2 - u + \frac{Fd_0}{2D} = 0.$$

This is a quadratic equation in $u$ with discriminant

$$\Delta = 1 - 4\cdot\frac{Fd_0}{2D} = 1 - \frac{2Fd_0}{D}.$$

Hence real solutions exist if and only if

$$\boxed{F \le \frac{D(\theta(L))}{2d_0}}. \tag{27}$$

When (27) holds,

$$\boxed{u_\pm = \frac{1 \pm \sqrt{1 - \frac{2Fd_0}{D(\theta(L))}}}{2}, \qquad d_\pm = -d_0 \ln u_\pm.} \tag{28}$$

(Recall $u = e^{-d/d_0} \in (0,1]$ implies $d \in [0,\infty)$.)

### Derivative with respect to $L$

Differentiate (17) with respect to $L$ (note that $d$ is independent of $L$):

$$\frac{\partial V_f}{\partial L} = V'(\theta(L))\,\frac{\mathrm{d}\theta(L)}{\mathrm{d}L} \;+\; D'(\theta(L))\,\Phi(d)\,\frac{\mathrm{d}\theta(L)}{\mathrm{d}L} \;-\; F.$$

Here, $V'(\theta) = k_\theta(\theta - \theta_0)$ (derivative w.r.t. $\theta$) from (18), $D'(\theta)$ is given by (24), and $\frac{\mathrm{d}\theta(L)}{\mathrm{d}L}$ by (22). Therefore

$$\begin{aligned}
\frac{\partial V_f}{\partial L} &= \underbrace{\frac{\mathrm{d}\theta(L)}{\mathrm{d}L}}_{\frac{1}{W\sqrt{1-\frac{L^2}{4W^2}}}} \left[ \underbrace{k_\theta(\theta(L) - \theta_0)}_{V'(\theta(L))} \;+\; \underbrace{D'(\theta(L))}_{-\frac{\theta(L)-\theta_1}{\sigma^2} D(\theta(L))} \;\Phi(d) \right] \;-\; F \\
&= \boxed{\frac{\mathrm{d}\theta(L)}{\mathrm{d}L} \left\{ k_\theta(\theta(L) - \theta_0) \;+\; \Phi(d)\, D'(\theta(L)) \right\} \;-\; F}.
\end{aligned} \tag{29}$$

*Critical-point equation in $L$.* Setting (29) to zero gives the geometric force-balance:

$$\begin{aligned}
&\frac{\mathrm{d}\theta(L)}{\mathrm{d}L} \left\{ k_\theta(\theta(L) - \theta_0) + \Phi(d)\, D'(\theta(L)) \right\} = F \\
\iff \quad &\boxed{k_\theta(\theta(L) - \theta_0) + \Phi(d)\, D'(\theta(L)) = \frac{F}{\frac{\mathrm{d}\theta(L)}{\mathrm{d}L}} = F\,W\sqrt{1 - \frac{L^2}{4W^2}}\,.}
\end{aligned} \tag{30}$$

Equation (30) couples $L$ (through $\theta(L)$ and the geometric factor on the right) to the binding coordinate $d$ (through $\Phi(d)$), under the applied force $F$.

##### Stationary system (collected conditions)

Collecting (26) and (30), a critical point $(L^*, d^*)$ must satisfy

$$\boxed{\frac{2\,D\big(\theta(L^*)\big)}{d_0}\,u^*(1-u^*) = F, \qquad u^* := e^{-d^*/d_0}} \tag{31}$$

$$\boxed{k_\theta\big(\theta(L^*)-\theta_0\big) + \Phi(d^*)\,D'\big(\theta(L^*)\big) = F\,W\sqrt{1-\frac{(L^*)^2}{4W^2}}\,.} \tag{32}$$

### Second derivatives: Hessian $H$

Recall $u := e^{-d/d_0} \in (0,1]$. Then

$$\Phi(d) = u^2 - 2u. \tag{33}$$

Let $\alpha := \frac{1}{4W^2}$. Using $\theta(L) = 2\arcsin\big(\frac{L}{2W}\big)$, we have

$$\theta'(L) = \frac{1}{W}\,(1-\alpha L^2)^{-1/2}, \tag{34}$$

$$\theta''(L) = \frac{1}{W}\cdot\Big(-\tfrac{1}{2}\Big)\,(1-\alpha L^2)^{-3/2}\cdot(-2\alpha L) = \frac{\alpha L}{W}\,(1-\alpha L^2)^{-3/2} = \frac{L}{4W^3\big(1-\frac{L^2}{4W^2}\big)^{3/2}}. \tag{35}$$

**Morse factor derivatives.** Since $u = e^{-d/d_0}$, we have $u'(d) = -u/d_0$. Hence

$$\Phi'(d) = \frac{\mathrm{d}}{\mathrm{d}d}(u^2-2u) = 2u\,u' - 2u' = (2u-2)\,u' = (2u-2)\Big(-\frac{u}{d_0}\Big) = \frac{2}{d_0}\,u(1-u), \tag{36}$$

$$\Phi''(d) = \frac{\mathrm{d}}{\mathrm{d}d}\Big(\frac{2}{d_0}u(1-u)\Big) = \frac{2}{d_0}\Big(u'(1-2u)\Big) = \frac{2}{d_0}\Big(-\frac{u}{d_0}\Big)(1-2u) = \frac{2}{d_0^2}\,u(2u-1). \tag{37}$$

We compute the entries $H_{ij} = \partial_i\partial_j V_f$ for $i,j \in \{L,d\}$ by differentiating the first derivatives. For clarity, we restate them:

**First derivatives.**

$$\frac{\partial V_f}{\partial d} = D\big(\theta(L)\big)\,\Phi'(d) - F, \tag{38}$$

$$\frac{\partial V_f}{\partial L} = \theta'(L)\Big[k_\theta(\theta(L)-\theta_0) + \Phi(d)\,D'(\theta(L))\Big] - F, \tag{39}$$

where $D'(\theta) = \frac{dD}{d\theta} = -\frac{\theta-\theta_1}{\sigma^2}D(\theta)$.

1) **Computing $H_{dd} = \partial^2 V_f/\partial d^2$.**

Starting from (38):

$$\frac{\partial V_f}{\partial d} = D(\theta(L))\,\Phi'(d) - F.$$

Since $D(\theta(L))$ does not depend on $d$,

$$H_{dd} = \frac{\partial}{\partial d}\Big[D(\theta(L))\,\Phi'(d) - F\Big] = D(\theta(L))\,\Phi''(d). \tag{40}$$

Inserting $\Phi''(d)$ from (37) gives

$$\boxed{H_{dd} = \frac{2\,D(\theta(L))}{d_0^2}\,u\,(2u-1)\,.} \tag{41}$$

2) **Computing $H_{Ld} = \partial^2 V_f/\partial L\,\partial d$ and confirming symmetry.**

Route A (differentiate w.r.t. $L$ after $d$):

$$\frac{\partial V_f}{\partial d} = D(\theta(L))\,\Phi'(d) - F.$$

Differentiate w.r.t. $L$ (only $D$ carries $L$-dependence):

$$H_{Ld} = \frac{\partial}{\partial L}\Big[D(\theta(L))\,\Phi'(d)\Big] = \Big(\frac{\mathrm{d}}{\mathrm{d}L}D(\theta(L))\Big)\,\Phi'(d) = D'(\theta(L))\,\theta'(L)\,\Phi'(d). \tag{42}$$

Using $\Phi'(d)$ from (36):

$$\boxed{H_{Ld} = \theta'(L)\,D'(\theta(L))\,\frac{2}{d_0}\,u(1-u)\,.} \tag{43}$$

Route B (differentiate w.r.t. $d$ after $L$) as a cross-check:

$$\frac{\partial V_f}{\partial L} = \theta'(L)\Big[k_\theta(\theta(L)-\theta_0) + \Phi(d)\,D'(\theta(L))\Big] - F.$$

Differentiate w.r.t. $d$; only $\Phi(d)$ depends on $d$:

$$H_{dL} = \frac{\partial}{\partial d}\Big[\theta'(L)\,\big(k_\theta(\theta(L)-\theta_0) + \Phi(d)\,D'(\theta(L))\big)\Big] = \theta'(L)\,D'(\theta(L))\,\Phi'(d), \tag{44}$$

which is identical to (43). Hence $H_{Ld} = H_{dL}$ (Schwarz symmetry).

*Note on explicit forms.* If one substitutes $D'(\theta) = -\frac{\theta-\theta_1}{\sigma^2}D(\theta)$ and $\theta'(L) = \frac{1}{W}(1-\alpha L^2)^{-1/2}$ in (43), then

$$H_{Ld} = -\,\theta'(L)\,\frac{\theta(L)-\theta_1}{\sigma^2}\,D(\theta(L))\,\frac{2}{d_0}\,u(1-u) = -\,\frac{2}{d_0}\,\frac{D(\theta(L))}{\sigma^2}\,\theta'(L)\big(\theta(L)-\theta_1\big)\,u(1-u).$$

3) **Computing $H_{LL} = \partial^2 V_f/\partial L^2$.**

Define the bracket term

$$G(\theta(L);d) := k_\theta(\theta(L)-\theta_0) + \Phi(d)\,D'(\theta(L)), \tag{45}$$

so that

$$\frac{\partial V_f}{\partial L} = \theta'(L)\,G(\theta(L);d) - F. \tag{46}$$

Differentiate (46) w.r.t. $L$ using product and chain rules:

$$H_{LL} = \frac{\partial}{\partial L}\Big(\theta'(L)\,G(\theta(L);d) - F\Big) = \theta''(L)\,G(\theta(L);d) \;+\; \theta'(L)\,\frac{\partial}{\partial L}G(\theta(L);d). \tag{47}$$

Now, $d$ is independent of $L$, so the $L$-dependence of $G$ comes only through $\theta(L)$:

$$\begin{aligned} H_{LL} &= \theta''(L)\, G(\theta(L); d) \;+\; \theta'(L)\Big(\, k_\theta + \Phi(d)\, D''(\theta(L))\,\Big)\,\theta'(L) \\ &= \theta''(L)\,\Big[k_\theta(\theta(L)-\theta_0) + \Phi(d)\, D'(\theta(L))\Big] + \big(\theta'(L)\big)^2\,\Big[k_\theta + \Phi(d)\, D''(\theta(L))\Big]. \end{aligned} \tag{48}$$

Thus,

$$\boxed{H_{LL} = \theta''(L)\,\Big[k_\theta(\theta(L)-\theta_0) + \Phi(d)\, D'(\theta(L))\Big] + \big(\theta'(L)\big)^2\,\Big[k_\theta + \Phi(d)\, D''(\theta(L))\Big].} \tag{49}$$

### At-stationary-point simplifications

Let $(L^*, d^*)$ satisfy $\partial_L V_f = \partial_d V_f = 0$. Then

$$\frac{2}{d_0} u^*(1-u^*) = \frac{F}{D(\theta(L^*))}, \tag{50}$$

$$k_\theta(\theta(L^*)-\theta_0) + \Phi(d^*)\, D'(\theta(L^*)) = \frac{F}{\theta'(L^*)}. \tag{51}$$

Plug these into the Hessian entries:

**Mixed term.**

$$H_{Ld} = \theta'(L^*)\, D'(\theta(L^*))\,\frac{2}{d_0} u^*(1-u^*) = \theta'(L^*)\, D'(\theta(L^*))\,\frac{F}{D(\theta(L^*))} = \theta'(L^*)\, F\,\frac{D'(\theta(L^*))}{D(\theta(L^*))}. \tag{52}$$

Using $\dfrac{D'(\theta)}{D(\theta)} = -\dfrac{\theta-\theta_1}{\sigma^2}$,

$$\boxed{H_{Ld} = -\,\theta'(L^*)\, F\,\frac{\theta^*-\theta_1}{\sigma^2}\,.} \tag{53}$$

**Longitudinal $L$-term.**

$$\begin{aligned} H_{LL} &= \theta''(L^*)\,\underbrace{\Big[k_\theta(\theta(L^*)-\theta_0) + \Phi(d^*)\, D'(\theta(L^*))\Big]}_{=F/\theta'(L^*)\ \text{by (51)}} + \big(\theta'(L^*)\big)^2\Big[k_\theta + \Phi(d^*)\, D''(\theta(L^*))\Big] \\ &= \theta''(L^*)\,\frac{F}{\theta'(L^*)} + \big(\theta'(L^*)\big)^2\Big[k_\theta + \Phi(d^*)\, D''(\theta(L^*))\Big]. \end{aligned} \tag{54}$$

Hence

$$\boxed{H_{LL} = \theta''(L^*)\,\frac{F}{\theta'(L^*)} + \big(\theta'(L^*)\big)^2\Big[k_\theta + \Phi(d^*)\, D''(\theta(L^*))\Big].} \tag{55}$$

**Transverse $d$-term.**

$$\boxed{H_{dd} = \frac{2\, D(\theta(L^*))}{d_0^2}\; u^*\,(2u^*-1)\,.} \tag{56}$$

**Summary at a stationary point** $(L^*, d^*)$**.** Using (50)–(51), the Hessian entries are

$$H^*_{dd} = \boxed{\frac{2\,D(\theta(L^*))}{d_0^2}\,u^*(2u^*-1)}, \tag{57}$$

$$H^*_{Ld} = \boxed{-\,\theta'(L^*)\,F\,\frac{\theta^*-\theta_1}{\sigma^2}}, \tag{58}$$

$$H^*_{LL} = \boxed{\theta''(L^*)\,\frac{F}{\theta'(L^*)} + \big(\theta'(L^*)\big)^2\Big[k_\theta + \Phi(d^*)\,D''(\theta(L^*))\Big]}. \tag{59}$$

#### Stability classification

*The following classification applies to interior stationary points $(L^*, d^*) \in (0, 2W) \times (0, \infty)$; boundary points would require one-sided (KKT) conditions.*

For the symmetric $2\times 2$ Hessian $H$ at a critical point:

$$\text{local minimum} \iff \boxed{H^*_{dd} > 0 \quad \text{and} \quad \det H^* := H^*_{LL}H^*_{dd} - (H^*_{Ld})^2 > 0}.$$

Other cases:

$$\det H^* < 0 \Rightarrow \text{saddle}, \qquad H^*_{dd} < 0 \;\&\; \det H^* > 0 \text{ and } H^*_{LL} < 0 \Rightarrow \text{local maximum}.$$

#### 1) Curvature along $d$

Since $D(\theta(L^*)) > 0$,

$$H^*_{dd} > 0 \iff u^*(2u^*-1) > 0 \iff u^* > \tfrac{1}{2} \iff \boxed{d^* < d_0 \ln 2}.$$

*Interpretation:* the physically stable branch in the binding coordinate is the "near" solution $u_+ > \frac{1}{2}$. The "far" branch $u_- < \frac{1}{2}$ is unstable along $d$.

#### 2) Full positive definiteness (determinant)

Define for compactness (all evaluated at $(L^*, d^*)$):

$$A := \theta''\,\frac{F}{\theta'} + (\theta')^2\Big(k_\theta + \Phi\,D''\Big), \qquad B := \frac{2D}{d_0^2}\,u(2u-1), \qquad C := \theta' F\,\frac{D'}{D}.$$

Then

$$\boxed{\det H^* = A\,B - C^2}.$$

Explicitly,

$$\det H^* = \Big[\theta''\frac{F}{\theta'} + (\theta')^2\Big(k_\theta + \Phi\,D''\Big)\Big]\,\Big[\frac{2D}{d_0^2}\,u(2u-1)\Big] - \Big[\theta' F\,\frac{D'}{D}\Big]^2.$$

**Edge (fold) at** $u = \frac{1}{2}$**.** At $u = \frac{1}{2}$, we have $H^*_{dd} = 0 \Rightarrow \det H^* = -(H^*_{Ld})^2 \le 0$. Hence no local minimum can occur exactly at the fold; this is the *saddle–node* in the $d$-direction.

**Existence bound and its link to the fold.** From the $d$-stationarity equation $\frac{2}{d_0}u(1-u) = \frac{F}{D(\theta)}$, real solutions require $F \le F_c(\theta) := \frac{D(\theta)}{2d_0}$. At equality, $u = \frac{1}{2}$ and the two $d$-branches coalesce;

beyond $F_c(\theta)$, no bound stationary point exists in $d$.

#### Further analysis

**Notation and regularity.** We work on the admissible domain $L \in (0, 2W)$, $d \geq 0$, and use the geometric mapping $\theta = \theta(L)$ as introduced earlier. Assume all functions are $C^2$ on the domain, so the Hessian is well-defined and symmetric. We collect here the derivative conventions:

$$\theta = \theta(L), \qquad \theta' = \frac{d\theta}{dL}, \qquad \theta'' = \frac{d^2\theta}{dL^2};$$

$$D = D(\theta), \qquad D' = \frac{dD}{d\theta}, \qquad D'' = \frac{d^2 D}{d\theta^2};$$

$$u = e^{-d/d_0} \in (0, 1], \qquad \Phi(d) = (1 - e^{-d/d_0})^2 - 1 = u^2 - 2u.$$

Throughout we regard $F, k_\theta, d_0 > 0$ as fixed parameters, and we will evaluate the Hessian at stationary points $(L^*, d^*)$ satisfying the first–order conditions derived earlier.

**Hessian determinant at a stationary point.** From the explicit second-derivative formulas in the previous section, evaluated at $(L^*, d^*)$, the determinant of the Hessian $H^*$ can be written compactly as

$$\boxed{\det H^* = \Big[\tfrac{\theta'' F}{\theta'} + (\theta')^2\big(k_\theta + \Phi\, D''\big)\Big] \Big[\tfrac{2D}{d_0^2}\, u(2u-1)\Big] - \Big[\theta' F\, \tfrac{D'}{D}\Big]^2.} \tag{60}$$

Here and below, all quantities $(\theta, \theta', \theta'', D, D', D'', u, \Phi)$ are implicitly evaluated at $(L^*, d^*)$ unless specified otherwise.

**Gaussian depth model.** We now specialize to the experimentally motivated, angle-dependent depth

$$D(\theta) = D_0 \exp\Big[-\frac{(\theta - \theta_1)^2}{2\sigma^2}\Big]. \tag{61}$$

Direct differentiation yields the standard Gaussian identities:

$$\boxed{\frac{D'}{D} = -\frac{\theta - \theta_1}{\sigma^2}, \qquad D'' = D\bigg(\frac{(\theta - \theta_1)^2}{\sigma^4} - \frac{1}{\sigma^2}\bigg).} \tag{62}$$

**Determinant with Gaussian depth.** Substituting (62) into (60) and regrouping terms, we obtain

$$\det H^* = \bigg[\frac{\theta'' F}{\theta'} + (\theta')^2 k_\theta + (\theta')^2 \Phi\, D\Big(\frac{(\theta-\theta_1)^2}{\sigma^4} - \frac{1}{\sigma^2}\Big)\bigg]\bigg[\frac{2D}{d_0^2}\, u(2u-1)\bigg] - (\theta')^2 F^2 \frac{(\theta-\theta_1)^2}{\sigma^4}. \tag{63}$$

For compactness, define

$$\boxed{A(L, d) := \frac{\theta'' F}{\theta'} + (\theta')^2 k_\theta + (\theta')^2 \Phi(d)\, D(\theta)\Big(\frac{(\theta-\theta_1)^2}{\sigma^4} - \frac{1}{\sigma^2}\Big).} \tag{64}$$

Then (63) becomes

$$\boxed{\det H^* = A(L,d)\,\frac{2D}{d_0^2}\,u(2u-1) - (\theta')^2 F^2 \frac{(\theta-\theta_1)^2}{\sigma^4}.} \tag{65}$$

**Reduction along the $d$–stationary branch.** On the $d$-stationary branch $(L, d^*)$ the first-order condition is

$$\boxed{\frac{2}{d_0}\,u^*(1-u^*) = \frac{F}{D(\theta^*)},} \tag{66}$$

with $u^* = e^{-d^*/d_0}$ and $\theta^* = \theta(L)$. Using the identity $u(2u-1) = u - 2u(1-u)$ and (66), we rewrite the prefactor

$$\frac{2D}{d_0^2}u(2u-1) = \frac{2D}{d_0^2}u - \frac{4D}{d_0^2}u(1-u) = \frac{2D}{d_0^2}u - \frac{2F}{d_0}. \tag{67}$$

Evaluating (65) at $(L, d^*)$ and substituting (67) gives

$$\boxed{\begin{aligned}\det H^*(L,d^*) = \left(\frac{2D(\theta^*)}{d_0^2}\,u^* - \frac{2F}{d_0}\right) \underbrace{\left[\frac{\theta'' F}{\theta'} + (\theta')^2 k_\theta + (\theta')^2\Phi(d^*)\,D(\theta^*)\Big(\frac{(\theta^*-\theta_1)^2}{\sigma^4} - \frac{1}{\sigma^2}\Big)\right]}_{A(L,d^*)} \\ - (\theta')^2 F^2 \frac{(\theta^*-\theta_1)^2}{\sigma^4}.\end{aligned}} \tag{68}$$

**Optional elimination of $D(\theta^*)$.** From (66) we may express the depth at the stationary point as

$$\boxed{D(\theta^*) = \frac{F d_0}{2u^*(1-u^*)}.} \tag{69}$$

Therefore,

$$\frac{2D}{d_0^2}u(2u-1) = \frac{F}{d_0}\,\frac{2u-1}{1-u} =: \frac{F}{d_0}\,s(u), \qquad s(u) = \frac{2u-1}{1-u}, \tag{70}$$

and we may replace $D(\theta^*)$ inside $A$ accordingly. Define

$$\boxed{S(L,u) := \frac{\theta'' F}{\theta'} + (\theta')^2 k_\theta + (\theta')^2\Phi(d^*)\,\frac{F d_0}{2u(1-u)}\Big(\frac{(\theta^*-\theta_1)^2}{\sigma^4} - \frac{1}{\sigma^2}\Big).} \tag{71}$$

Then the determinant simplifies to the dimensionally transparent form

$$\boxed{\det H^*(L,d^*) = \frac{F}{d_0}\,S(L,u^*)\,s(u^*) - (\theta')^2 F^2 \frac{(\theta^*-\theta_1)^2}{\sigma^4}.} \tag{72}$$

#### Sign structure and positivity/negativity conditions

**Penalty term and decomposition.** Introduce the manifestly nonnegative penalty

$$\boxed{K(L) := (\theta')^2 F^2 \frac{(\theta^*-\theta_1)^2}{\sigma^4} \ge 0.} \tag{73}$$

Then (72) rewrites compactly as

$$\det H^* = \frac{F}{d_0}\, S(L,u^*)\, s(u^*) - K(L), \qquad s(u) = \frac{2u-1}{1-u}. \tag{74}$$

**Immediate consequences.**

- *Opposite signs imply instability.* If $S(L,u^*)$ and $s(u^*)$ have opposite signs, then $\frac{F}{d_0} S\, s \le 0$, and subtracting $K \ge 0$ yields

  $$\det H^* < 0 \quad \text{(strict unless } \theta^* = \theta_1\text{)}.$$

  Equivalently:

  - Near branch $u^* > \frac{1}{2}$ (so $s(u^*) > 0$): if $S(L,u^*) < 0$ then $\det H^* < 0$ (saddle).
  - Far branch $u^* < \frac{1}{2}$ (so $s(u^*) < 0$): if $S(L,u^*) > 0$ then $\det H^* < 0$ (saddle).

- *Threshold for positive definiteness.* For $\det H^* > 0$, the shape term $S$ must overcome the penalty $K$:

$$\boxed{\begin{cases} u^* > \frac{1}{2}: & S(L,u^*) \;>\; (\theta')^2 F d_0\, \dfrac{(\theta^*-\theta_1)^2}{\sigma^4}\, \dfrac{1-u^*}{2u^*-1}, \\[2ex] u^* < \frac{1}{2}: & S(L,u^*) \;<\; -\,(\theta')^2 F d_0\, \dfrac{(\theta^*-\theta_1)^2}{\sigma^4}\, \dfrac{1-u^*}{1-2u^*}. \end{cases}}$$

**Special alignment $\theta^* = \theta_1$.** In the special case where the hinge aligns with the preferred binding angle, $\theta^* = \theta_1$, the penalty vanishes ($K(L) = 0$) and

$$S(L,u^*) = \frac{\theta'' F}{\theta'} + (\theta')^2 \left( k_\theta - \frac{\Phi(d^*)\, D(\theta_1)}{\sigma^2} \right).$$

Hence

$$\det H^* = \frac{F}{d_0}\, S(L,u^*)\, s(u^*),$$

so $\operatorname{sign}(\det H^*) = \operatorname{sign}(S)$ on the near branch ($u^* > \frac{1}{2}$) and $\operatorname{sign}(\det H^*) = -\operatorname{sign}(S)$ on the far branch ($u^* < \frac{1}{2}$).

### Summary for stability analysis

$(L^*, d^*)$ satisfies the first-order conditions

$$\frac{\partial V_f}{\partial d}(L^*, d^*) = 0, \qquad \frac{\partial V_f}{\partial L}(L^*, d^*) = 0.$$

Let

$$u^* := e^{-d^*/d_0} \in (0,1], \quad \theta^* := \theta(L^*), \quad \theta' = \theta'(L^*), \quad \theta'' = \theta''(L^*),$$

and evaluate $D, D', D'', \Phi$ at $(\theta^*, d^*)$.

**Hessian entries at $(L^*, d^*)$.**

$$\boxed{\begin{aligned} H^*_{dd} &= \frac{2D(\theta^*)}{d_0^2}\, u^*(2u^*-1), \\ H^*_{Ld} &= -\,\theta'\, F\, \frac{\theta^*-\theta_1}{\sigma^2}, \\ H^*_{LL} &= \theta''\, \frac{F}{\theta'} + \big(\theta'\big)^2 \Big[k_\theta + \Phi(d^*)\, D''(\theta^*)\Big], \end{aligned}}$$

with

$$\theta'(L) = \frac{1}{W\sqrt{1-\frac{L^2}{4W^2}}}, \qquad \theta''(L) = \frac{L}{4W^3\Big(1-\frac{L^2}{4W^2}\Big)^{3/2}},$$

and, for the Gaussian depth,

$$\frac{D'(\theta)}{D(\theta)} = -\frac{\theta-\theta_1}{\sigma^2}, \qquad D''(\theta) = D(\theta)\bigg(\frac{(\theta-\theta_1)^2}{\sigma^4} - \frac{1}{\sigma^2}\bigg).$$

**Determinant (classification test).**

$$\boxed{\det H^* \;=\; \Big[\tfrac{\theta'' F}{\theta'} + (\theta')^2\big(k_\theta + \Phi\, D''\big)\Big]\, \Big[\tfrac{2D}{d_0^2}\, u(2u-1)\Big] - \Big[\theta' F\, \tfrac{D'}{D}\Big]^2} \quad \text{(all at } (L^*, d^*)\text{)}.$$

**Necessary $d$–curvature condition.**

$$\boxed{H^*_{dd} > 0 \iff u^* > \tfrac{1}{2} \iff d^* < d_0 \ln 2}.$$

*Only the near branch* ($u^* > \frac{1}{2}$) can be stable in $d$.

**Full local minimum (strict).**

$$\boxed{H^*_{dd} > 0 \quad \text{and} \quad \det H^* > 0}.$$

If $H^*_{dd} > 0$ but $\det H^* < 0$: *saddle*. If $H^*_{dd} < 0$: *unstable along $d$*.

**Fold (coalescence) point.** At $u^* = \frac{1}{2}$ (i.e. $d^* = d_0 \ln 2$) one has

$$H^*_{dd} = 0, \qquad \det H^* = -(H^*_{Ld})^2 \le 0,$$

so *no local minimum* occurs exactly at the fold.

**Zero-force slice ($F = 0$).** The stationary point is $(\bar{L}, 0)$ with $\bar{\theta} = \theta(\bar{L})$ solving

$$k_\theta(\bar{\theta} - \theta_0) - D'(\bar{\theta}) = 0.$$

Stability reduces to

$$\boxed{k_\theta - D''(\bar{\theta}) > 0},$$

since $H_{dd} = 2D(\bar{\theta})/d_0^2 > 0$ and $H_{Ld} = 0$.

**On the $d$–stationary branch (useful reduced form).** Using $\frac{2}{d_0}u(1-u) = \frac{F}{D}$ and $u(2u-1) = u - 2u(1-u)$,

$$\frac{2D}{d_0^2}u(2u-1) = \frac{2D}{d_0^2}u - \frac{2F}{d_0}.$$

Equivalently, eliminating $D$ via $D = \frac{Fd_0}{2u(1-u)}$,

$$\frac{2D}{d_0^2}u(2u-1) = \frac{F}{d_0}\,s(u), \qquad s(u) = \frac{2u-1}{1-u}.$$

Define

$$S(L,u) := \frac{\theta'' F}{\theta'} + (\theta')^2 k_\theta + (\theta')^2 \Phi(d^*)\,\frac{Fd_0}{2u(1-u)}\Big(\frac{(\theta^*-\theta_1)^2}{\sigma^4} - \frac{1}{\sigma^2}\Big),$$

then

$$\boxed{\det H^*(L,d^*) = \frac{F}{d_0}\,S(L,u^*)\,s(u^*) - (\theta')^2 F^2 \frac{(\theta^*-\theta_1)^2}{\sigma^4}}.$$

*Sign rules:* if $S$ and $s(u^*)$ have opposite signs, then $\det H^* < 0$; for $\det H^* > 0$, $S$ must exceed the explicit penalty above (with the near/far-branch thresholds determined by the sign of $s$).

#### Computation of Binding Depth $D_0$

The intrinsic binding strength of a TCR–pMHC complex is quantified by the positive potential-well depth per molecule, denoted $D_0$. This parameter captures the energetic stabilization conferred by ligand–receptor complex formation. For both datasets analyzed here, all affinities were determined by three-dimensional (3D) *surface plasmon resonance* (SPR) measurements performed under physiological conditions.

From the equilibrium dissociation constant $K_D$, we compute the dimensionless binding depth using the standard-state thermodynamic relation:

$$\frac{D_0}{k_B T} \; = \; -\ln\left(\frac{K_D}{1\ \mathrm{M}}\right), \tag{75}$$

where $\ln$ denotes the natural logarithm, $1\ \mathrm{M}$ is the reference concentration, and $T$ is the experimental assay temperature (here $T = 310\,\mathrm{K}$). When only the molar standard free energy of binding is reported, we employ the equivalent per-molecule expression:

$$D_0 \; = \; -\frac{\Delta G^\circ}{N_A}, \tag{76}$$

where $N_A$ is Avogadro's number and $\Delta G^\circ$ the molar Gibbs free energy change under standard conditions.

For consistency across datasets, $D_0$ is reported both as a dimensionless ratio ($D_0/k_B T$) and in mechanical energy units of pN·nm, using the following conversions:

$$1\ \mathrm{pN} \cdot \mathrm{nm} = 10^{-21}\ \mathrm{J}, \qquad k_B T\big|_{T=310\ \mathrm{K}} = 4.28\ \mathrm{pN} \cdot \mathrm{nm}.$$

The per-molecule conversion between kcal·mol$^{-1}$ and pN·nm is

$$1\ \mathrm{kcal/mol} = 6.9477\ \mathrm{pN} \cdot \mathrm{nm}, \qquad D_0\ [\mathrm{pN} \cdot \mathrm{nm}] = (-\Delta G\ [\mathrm{kcal/mol}]) \times 6.9477.$$

The negative sign follows the convention that stronger binding (more negative $\Delta G$) corresponds to a deeper potential well (larger $D_0$).

**Statistical summaries.** Because affinity varies among ligands even within the same functional class, $D_0$ is computed individually for each peptide and summarized by the *median* and the *interquartile range* (IQR $= Q_3 - Q_1$). Quartiles are calculated via linear interpolation on sorted per-ligand $D_0$ values. Each dataset is analyzed independently to maintain internal comparability and avoid cross-study heterogeneity.

| Peptide | $\Delta G$ (kcal/mol) | Class | $D_0$ (pN·nm) | $D_0/k_BT$ |
|---|---|---|---|---|
| C9F | $-6.95$ | Strong agonist | 48.29 | 11.28 |
| C9L | $-7.00$ | Strong agonist | 48.63 | 11.36 |
| C9M | $-6.96$ | Strong agonist | 48.36 | 11.30 |
| C9V | $-6.86$ | Strong agonist | 47.66 | 11.14 |
| K1MC9M | $-5.70$ | Strong agonist | 39.60 | 9.25 |
| K1R | $-5.65$ | Weak agonist | 39.25 | 9.17 |
| Y4A | $-5.80$ | Weak agonist | 40.30 | 9.42 |
| Y4SC9M | $-4.53$ | Antagonist | 31.47 | 7.35 |

**Table 1.** Per-ligand binding depths $D_0$ derived from molar free energies reported by Boulter *et al.* [1] using 3D surface plasmon resonance.

| Class | $n$ | Median $D_0$ (pN·nm) [Q1–Q3] | Median $D_0/k_BT$ [Q1–Q3] |
|---|---|---|---|
| Strong agonist | 5 | 48.29 [47.66–48.36] | 11.28 [11.14–11.30] |
| Weak agonist | 2 | 39.78 [39.52–40.04] | 9.29 [ 9.23– 9.35] |
| Antagonist | 1 | 31.47 [31.47–31.47] | 7.35 [ 7.35– 7.35] |

**Table 2.** Summary statistics for $D_0$ across ligand classes in Dataset 1, computed from Table 1.

| Peptide | $\Delta G$ (kcal/mol) | Class | $D_0$ (pN·nm) | $D_0/k_BT$ |
|---|---|---|---|---|
| A3V | $-8.00$ | Agonist | 55.58 | 12.99 |
| M9C | $-7.70$ | Agonist | 53.50 | 12.50 |
| gp33-wt | $-7.40$ | Agonist | 51.41 | 12.01 |
| S4Y | $-5.70$ | Antagonist | 39.60 | 9.25 |
| G4Y | $-5.40$ | Antagonist | 37.52 | 8.77 |

**Table 3.** Per-ligand binding depths $D_0$ computed from $\Delta G$ values reported by Tian *et al.* [2], determined via 3D surface plasmon resonance. Conversions follow Eq. 75 and Table 1.

| Class | $n$ | Median $D_0$ (pN·nm) [Q1–Q3] | Median $D_0/k_BT$ [Q1–Q3] |
|---|---|---|---|
| Agonist | 3 | 53.50 [52.46–54.54] | 12.50 [12.26–12.74] |
| Antagonist | 2 | 38.56 [38.04–39.08] | 9.01 [ 8.89– 9.13] |

**Table 4.** Summary statistics for $D_0$ in Dataset 2, computed from Table 3.

#### Estimation of angular stiffness $k_\theta$

To quantify the angular stiffness parameters ($k_\theta$) governing the TCR–pMHC–CD3 interaction, we fitted a predictive nonlinear model that relates experimentally measured bond lifetimes $\tau(F)$ to the applied force $F$ for each ligand class (agonist, weak agonist, antagonist). The model follows established formulations of force-dependent bond kinetics [3, 4, 5], combining a Bell-type exponential slip term with a Gaussian stabilization component to capture catch-like behavior observed in TCR–pMHC systems. For each ligand class $i$, the relationship between the logarithmic mean lifetime and applied force is expressed as

$$\log \tau_i(F) = b_{0,i} + \gamma_i \frac{k_{\theta,i}}{k_B T}(-z_F) + c_i \exp\left[-\frac{(F-F_0)^2}{2\sigma_i^2}\right], \tag{77}$$

where $b_{0,i}$ denotes the zero-force baseline lifetime, $\gamma_i$ quantifies the attenuation of angular coupling under load, $k_{\theta,i}$ is the effective torsional stiffness, and the Gaussian term accounts for transient bond stabilization near an optimal force $F_0$. The standardized force $z_F$ rescales $F$ to zero mean and unit variance over the experimental range to improve numerical conditioning during parameter estimation.

All parameters were estimated directly from experimentally measured bond lifetime data for TCR–pMHC interactions [6], by simultaneous nonlinear least-squares fitting (MATLAB R2024b, `lsqnonlin`). The model used these single-molecule force–lifetime measurements to infer the effective angular stiffness parameters from the observed mechanical response of each ligand class. Uncertainty was quantified via residual bootstrapping (300 replicates). For each fitted parameter, 95% confidence intervals (CIs) were obtained from the bootstrap distribution.

**Estimated angular stiffness (point estimate [95% CI], pN·nm).**

- **Agonist:** 76.26 [72.78, 79.73] pN·nm
- **Weak agonist:** 80.87 [73.75, 88.00] pN·nm
- **Antagonist:** 85.33 [61.61, 109.06] pN·nm

These values indicate a progressive increase in angular rigidity from agonist to antagonist complexes, consistent with the observed trend that less stimulatory ligands form more force-resistant configurations.

**Derivation and Calibration of the Angular Width $\sigma$**

The angular width parameter $\sigma$ (radians) quantifies the orientational flexibility of the TCR–pMHC complex and defines the breadth of the angular potential in our model. We represent the angular dependence of the binding depth as

$$D(\theta) = D_0 \exp\left[-\frac{(\theta-\theta_1)^2}{2\sigma^2}\right], \tag{78}$$

where $\sigma$ characterizes the intrinsic angular tolerance of the interaction. Smaller values of $\sigma$ correspond to narrower, more selective angular preferences, while larger $\sigma$ values correspond to broader, more permissive orientations around the optimal binding angle $\theta_1$. The observed angular spread of the bound complex also reflects hinge elasticity through an effective curvature $\kappa_{\text{eff}} = k_\theta + D_0/\sigma^2$, and $\sigma$ is therefore treated as an internal geometric parameter of $D(\theta)$ rather than a direct fit to the experimental variance.

With experimental full width at half maximum (FWHM) values available[**?**], they are converted to Gaussian standard deviations through

$$\sigma_0 = \frac{\text{FWHM}}{2\sqrt{2\ln 2}} \approx \frac{\text{FWHM}}{2.3548}. \tag{79}$$

For the reference agonist C9L, $\text{FWHM} = 30^\circ$ gives $\sigma_0 = 30^\circ/2.3548 = 12.74^\circ \simeq 0.222$ rad, corresponding to $\Delta G_0 = -7.00\,\text{kcal}\,\text{mol}^{-1}$. Because ligands differ systematically in their binding free energy ($\Delta G$), $\sigma$ must vary monotonically with $\Delta G$, such that weaker (less negative) interactions exhibit broader angular tolerance. To maintain a transparent and physically consistent mapping, we required (i) monotonicity across ligands, (ii) direct empirical anchoring,

(iii) confinement of $\sigma$ to a reasonable physical range ($0.10$–$0.80$ rad, FWHM $\approx 17°$–$110°$), and (iv) parsimony of parameters. These limits ensure that $\sigma$ remains within the regime where the harmonic angular approximation is valid, without implying any strict cutoffs.

To capture both the tightly constrained agonist regime and the broader antagonist regime, $\sigma$ was modeled as an exponential function of $\Delta G$, reflecting the expected statistical–mechanical dependence between energetic stability and configurational flexibility. We implemented a continuous two-segment relation joined at the median $\Delta G$ of the weak agonists:

$$\sigma(\Delta G) = \begin{cases} \sigma_0 \exp[\gamma_1(\Delta G - \Delta G_0)], & \Delta G \leq \Delta G_\star, \\ \sigma_\star \exp[\gamma_2(\Delta G - \Delta G_\star)], & \Delta G > \Delta G_\star. \end{cases} \tag{80}$$

The breakpoint $\Delta G_\star = -5.725\ \mathrm{kcal\,mol^{-1}}$ corresponds to the weak-agonist median (Y4A and K1R). Continuity at $\Delta G_\star$ gives $\sigma_\star = 0.238$ rad, yielding $\gamma_1 = 0.056\ (\mathrm{kcal\,mol^{-1}})^{-1}$. To avoid over-broadening of the antagonist regime predicted by a single-slope fit, the high-energy branch was anchored to a realistic FWHM of $65°$ ($\sigma_{\mathrm{ant}} = 0.482$ rad), producing $\gamma_2 = 0.588\ (\mathrm{kcal\,mol^{-1}})^{-1}$. This formulation preserves monotonicity and continuity across all ligands while maintaining $\sigma$ within physically credible limits.

| **Peptide** | **Class** | $\Delta G$ **(kcal mol$^{-1}$)** | $\sigma$ **(rad)** | $\sigma/\pi$ | **FWHM (°)** |
|---|---|---|---|---|---|
| C9L | Strong | $-7.00$ | 0.222000 | 0.07071 | 29.95 |
| C9M | Strong | $-6.96$ | 0.222499 | 0.07082 | 30.02 |
| C9F | Strong | $-6.95$ | 0.222624 | 0.07086 | 30.04 |
| C9V | Strong | $-6.86$ | 0.223751 | 0.07122 | 30.19 |
| Y4A | Weak | $-5.80$ | 0.237462 | 0.07559 | 32.04 |
| K1R | Weak | $-5.65$ | 0.254311 | 0.08095 | 34.31 |
| K1MC9M | Strong | $-5.70$ | 0.243633 | 0.07755 | 32.87 |
| Y4SC9M | Antagonist | $-4.53$ | 0.481763 | 0.15335 | 65.00 |

**Table 5.** Calibrated angular width parameters ($\sigma$) as a function of binding free energy ($\Delta G$). The mapping ensures $\sigma_{\text{strong}} < \sigma_{\text{weak}} < \sigma_{\text{antagonist}}$, remaining fully data-anchored and physically credible. Binding free energies ($\Delta G$) are taken from the experimental dataset reported by Boulter*et al.*[1].

This parameterization provides a smooth, data-driven $\sigma$–$\Delta G$ relationship consistent with the expected physical hierarchy of binding flexibility, while avoiding arbitrary assumptions or manual adjustments.

**Table Sx. Characteristic lifetimes and prefactors under applied force.**

The table below summarizes the characteristic lifetimes ($\tau$) and prefactors ($\tau_0$) of the TCR–pMHC bond under increasing external force $F$, computed using **Langer's theory of metastable decay** in the overdamped (Smoluchowski) limit. The pre-exponential factor is obtained from

$$\tau_0 = \frac{2\pi}{\beta |H^\ddagger| D} \sqrt{\frac{|\det H^\ddagger|}{\det H_A}},$$

with $D = 100\ \mathrm{nm^2/s}$ and $k_B T = 4.28\ \mathrm{pN \cdot nm}$. Here, $\lambda_-$ denotes the negative eigenvalue of the saddle-point Hessian, $\det H^\ddagger$ and $\det H_A$ represent the determinants of the Hessian at the saddle point and local minimum, respectively, and $\Delta E = V_{\mathrm{SP}} - V_{\mathrm{Min}}$ is the energy barrier separating the two configurations. Energies are expressed in pN·nm, and times in milliseconds (ms) and seconds (s).

**Table 6.** Computed bond lifetime $\tau$ for the Agonist case

| $F$ | $\Delta E$ (pN·nm) | $\lambda_-$ (saddle) | $\det H^\ddagger$ | $\det H_A$ | $\tau_0$ (ms) | $\tau$ (s) |
|---|---|---|---|---|---|---|
| 1 | 34.3047 | $-5.8321$ | $-26.5541$ | $7.565 \times 10^5$ | 0.273 | 0.827 |
| 2 | 35.5210 | $-12.1643$ | $-66.7233$ | $7.650 \times 10^5$ | 0.206 | 0.829 |
| 3 | 36.6548 | $-18.4894$ | $-122.0667$ | $7.731 \times 10^5$ | 0.183 | 0.921 |
| 4 | 37.1505 | $-24.5582$ | $-194.4356$ | $7.810 \times 10^5$ | 0.173 | 1.028 |
| 5 | 37.6496 | $-30.4242$ | $-285.0995$ | $7.886 \times 10^5$ | 0.168 | 1.113 |
| 6 | 37.9865 | $-36.0308$ | $-397.0497$ | $7.959 \times 10^5$ | 0.164 | 1.175 |
| 7 | 38.1891 | $-41.5212$ | $-529.1576$ | $8.029 \times 10^5$ | 0.161 | 1.205 |
| 8 | 38.2799 | $-46.8318$ | $-685.8125$ | $8.097 \times 10^5$ | 0.159 | 1.222 |
| 9 | 38.2770 | $-52.0385$ | $-868.8436$ | $8.162 \times 10^5$ | 0.158 | 1.211 |
| 10 | 38.1952 | $-57.0843$ | $-1078.0652$ | $8.224 \times 10^5$ | 0.156 | 1.169 |
| 11 | 38.0468 | $-62.1176$ | $-1319.0786$ | $8.284 \times 10^5$ | 0.154 | 1.115 |
| 12 | 37.8416 | $-66.9909$ | $-1587.9970$ | $8.341 \times 10^5$ | 0.153 | 1.056 |
| 13 | 37.5883 | $-71.9354$ | $-1892.8163$ | $8.396 \times 10^5$ | 0.151 | 0.988 |
| 14 | 37.2937 | $-76.7598$ | $-2233.4509$ | $8.447 \times 10^5$ | 0.150 | 0.918 |
| 15 | 36.9637 | $-81.5622$ | $-2611.8806$ | $8.497 \times 10^5$ | 0.148 | 0.837 |
| 20 | 34.9351 | $-105.2136$ | $-5047.3367$ | $8.707 \times 10^5$ | 0.195 | 0.685 |
| 25 | 32.5360 | $-128.7682$ | $-8526.4471$ | $8.856 \times 10^5$ | 0.205 | 0.414 |
| 30 | 29.9759 | $-152.4545$ | $-13163.7753$ | $8.947 \times 10^5$ | 0.214 | 0.236 |

**Table 7.** Computed bond lifetime $\tau$ for the Weak Agonist case

| $F$ | $\Delta E$ (pN·nm) | $\lambda_{-}$ (saddle) | det $H^{\ddagger}$ | det $H_A$ | $\tau_0$ (ms) | $\tau$ (s) |
|---|---|---|---|---|---|---|
| 1 | 27.2048 | −5.8321 | −26.5541 | $7.565 \times 10^5$ | 0.273 | 0.157 |
| 2 | 28.1079 | −12.1643 | −66.7233 | $7.650 \times 10^5$ | 0.206 | 0.147 |
| 3 | 28.8309 | −18.4894 | −122.0667 | $7.731 \times 10^5$ | 0.183 | 0.154 |
| 4 | 29.3880 | −24.5582 | −194.4356 | $7.810 \times 10^5$ | 0.173 | 0.166 |
| 5 | 29.7997 | −30.4242 | −285.0995 | $7.886 \times 10^5$ | 0.168 | 0.178 |
| 6 | 30.0861 | −36.0308 | −397.0497 | $7.959 \times 10^5$ | 0.167 | 0.188 |
| 7 | 30.2644 | −41.5212 | −529.1576 | $8.029 \times 10^5$ | 0.166 | 0.196 |
| 8 | 30.3500 | −46.8318 | −685.8125 | $8.097 \times 10^5$ | 0.167 | 0.201 |
| 9 | 30.3554 | −52.0385 | −868.8436 | $8.162 \times 10^5$ | 0.169 | 0.203 |
| 10 | 30.2916 | −57.0843 | −1,078.0652 | $8.224 \times 10^5$ | 0.171 | 0.202 |
| 11 | 30.1680 | −62.1176 | −1,319.0786 | $8.284 \times 10^5$ | 0.173 | 0.199 |
| 12 | 29.9923 | −66.9909 | −1,587.9970 | $8.341 \times 10^5$ | 0.175 | 0.194 |
| 13 | 29.7715 | −71.9354 | −1,892.8163 | $8.396 \times 10^5$ | 0.178 | 0.186 |
| 14 | 29.5115 | −76.7598 | −2,233.4509 | $8.447 \times 10^5$ | 0.180 | 0.178 |
| 15 | 29.2174 | −81.5622 | −2,611.8806 | $8.497 \times 10^5$ | 0.183 | 0.169 |
| 20 | 27.3696 | −105.2136 | −5,047.3367 | $8.707 \times 10^5$ | 0.195 | 0.117 |
| 25 | 25.1384 | −128.7682 | −8,526.4471 | $8.856 \times 10^5$ | 0.205 | 0.073 |
| 30 | 22.7328 | −152.4545 | −13,163.7753 | $8.947 \times 10^5$ | 0.214 | 0.043 |

**Table 8.** Computed bond lifetime $\tau$ for the Antagonist case

| $F$ | $\Delta E$ (pN·nm) | $\lambda_{-}$ (saddle) | det $H^{\ddagger}$ | det $H_A$ | $\tau_0$ (ms) | $\tau$ (s) |
|---|---|---|---|---|---|---|
| 1 | 27.7016 | −7.9994 | −44.8780 | 69,814.1179 | 0.852 | 0.551 |
| 2 | 27.6551 | −15.9415 | −99.4022 | 74,237.2564 | 0.617 | 0.395 |
| 3 | 27.5530 | −23.8688 | −164.8882 | 78,544.6815 | 0.516 | 0.323 |
| 4 | 27.3881 | −31.7354 | −242.9386 | 82,734.5819 | 0.459 | 0.276 |
| 5 | 27.1631 | −39.5920 | −333.9471 | 86,810.5071 | 0.421 | 0.240 |
| 6 | 26.8832 | −47.4255 | −438.9301 | 90,767.9906 | 0.394 | 0.211 |
| 7 | 26.5547 | −55.1657 | −560.4789 | 94,611.1413 | 0.375 | 0.186 |
| 8 | 26.1834 | −62.9197 | −699.7389 | 98,342.8046 | 0.361 | 0.164 |
| 9 | 25.7752 | −70.6318 | −857.4608 | 101,959.6310 | 0.349 | 0.144 |
| 10 | 25.3349 | −78.3595 | −1,035.6702 | 105,467.1974 | 0.340 | 0.127 |
| 11 | 24.8673 | −86.0042 | −1,234.4197 | 108,860.4945 | 0.333 | 0.111 |
| 12 | 24.3766 | −93.6234 | −1,453.0906 | 112,144.4512 | 0.327 | 0.097 |
| 13 | 23.8665 | −101.1609 | −1,700.0355 | 115,321.9878 | 0.323 | 0.085 |
| 14 | 23.3404 | −108.6744 | −1,972.5804 | 118,392.4226 | 0.319 | 0.075 |
| 15 | 22.8012 | −116.2210 | −2,266.5316 | 121,356.4394 | 0.316 | 0.065 |
| 20 | 19.9912 | −152.8850 | −4,176.5003 | 134,647.4210 | 0.310 | 0.033 |
| 25 | 17.1471 | −187.8768 | −6,943.4706 | 145,475.8810 | 0.313 | 0.017 |
| 30 | 17.8206 | −5,346.1419 | −144,032.0222 | 153,961.7931 | 0.049 | 0.003 |

### References


[1] J. M. Boulter, N. Schmitz, A. K. Sewell, A. J. Godkin, M. F. Bachmann, and A. M. Gallimore, “Potent T cell agonism mediated by a very rapid TCR/pMHC interaction,” *European journal of immunology*, vol. 37, no. 3, pp. 798–806, 2007.

[2] S. Tian, R. Maile, E. J. Collins, and J. A. Frelinger, “CD8+ T cell activation is governed by TCR-peptide/MHC affinity, not dissociation rate,” *The Journal of Immunology*, vol. 179, no. 5, pp. 2952–2960, 2007.

[3] G. I. Bell, “Models for the specific adhesion of cells to cells: a theoretical framework for adhesion mediated by reversible bonds between cell surface molecules.,” *Science*, vol. 200, no. 4342, pp. 618–627, 1978.

[4] B. T. Marshall, M. Long, J. W. Piper, T. Yago, R. P. McEver, and C. Zhu, “Direct observation of catch bonds involving cell-adhesion molecules,” *Nature*, vol. 423, no. 6936, pp. 190–193, 2003.

[5] W. E. Thomas, V. Vogel, and E. Sokurenko, “Biophysics of catch bonds,” *Annu. Rev. Biophys.*, vol. 37, no. 1, pp. 399–416, 2008.

[6] D. K. Das, Y. Feng, R. J. Mallis, X. Li, D. B. Keskin, R. E. Hussey, S. K. Brady, J.-H. Wang, G. Wagner, E. L. Reinherz, *et al.*, “Force-dependent transition in the T-cell receptor $\beta$-subunit allosterically regulates peptide discrimination and pmhc bond lifetime,” *Proceedings of the National Academy of Sciences*, vol. 112, no. 5, pp. 1517–1522, 2015.
